# Anti-angiogenic effects of VEGF stimulation on endothelium deficient in phosphoinositide recycling

**DOI:** 10.1101/402362

**Authors:** Amber N. Stratman, Olivia M. Farrelly, Constantinos M. Mikelis, Mayumi F. Miller, Zhiyong Wang, Van N. Pham, Andrew E. Davis, Margaret C. Burns, Sofia A. Pezoa, Daniel Castranova, Joseph J. Yano, Tina M. Kilts, George E. Davis, J. Silvio Gutkind, Brant M. Weinstein

## Abstract

Anti-angiogenic therapies have generated significant interest for their potential to combat tumor growth (*1-6*). However, the ability of tumors to overproduce pro-angiogenic ligands and overcome targeted inhibitory therapies has hampered this approach (*7, 8*). A novel way to circumvent this problem might be to target the resynthesis of critical substrates consumed during intracellular transduction of pro-angiogenic signals in endothelial cells, thus harnessing the tumor’s own production of excess stimulatory ligands to deplete adjacent host endothelial cells of the capacity to respond to these signals (*9-12*). Here we show using zebrafish and human endothelial cells *in vitro* that endothelial cells deficient in *CDP-diacylglycerol synthase 2* are uniquely sensitive to increased VEGF stimulation due to a reduced capacity to re-synthesize phosphoinositides, including phosphatidylinositol 4,5-bisphosphate (PIP2) a key substrate for VEGF signal transduction, resulting in VEGF-exacerbated defects in angiogenesis and angiogenic signaling (*9-22*). Using murine tumor allograft models (*23*) we show that either systemic or endothelial cell specific suppression of phosphoinositide recycling results in reduced tumor growth and reduced tumor angiogenesis. Our results suggest that inhibition of phosphoinositide recycling may provide a useful anti-angiogenic approach, and highlights the general potential of targeting the resynthesis of rate limiting signaling substrates as a valuable therapeutic strategy.

**SUMMARY STATEMENT:** Targeting phosphoinositide recycling during tumor angiogenesis provides a potentially uniquely effective anti-cancer therapy.

---

Previously, we reported the discovery of a zebrafish mutant in the *CDP-diacylglycerol synthase (cds2)* gene with defects in angiogenesis (*9*). CDS activity is required for resynthesis of phosphoinositides, including phosphatidylinositol-(4,5)-bisphosphate, or PIP2 (*9-11, 24*) (**Fig. 1A**). Phosphoinositides are utilized as “second messengers” or substrates for a wide variety of different intracellular signaling pathways. In endothelial cells (EC), PIP2 serves as a key substrate for both phospholipase C-gamma 1 (PLCg1) and phosphoinositide 3-kinase (PI3K) dependent signaling downstream from the VEGF-A receptor VEGFR2 (*12, 15-18, 20-22*) (**Fig. 1B**). To regenerate PIP2 consumed during VEGF/PLCg1 signaling, diacylglycerol (DAG) is recycled to CDP-diacyglycerol (CDP-DAG) through the activity of the two vertebrate CDP-diacyglycerol synthase genes, CDS1 and CDS2 (*9-22*). Loss or knockdown of either one of the two CDS genes in endothelial cells *in vitro* or in zebrafish embryos *in vivo* results in reduced angiogenesis that can be rescued by artificial elevation of PIP2 levels (*9*). Importantly, vascular defects are observed in *cds2*^*y54*^ mutant zebrafish embryos in the absence of other obvious developmental abnormalities, suggesting ECs are more sensitive to partial reduction in phosphoinositide recycling capacity than other cells and tissues.

**Figure 1.**
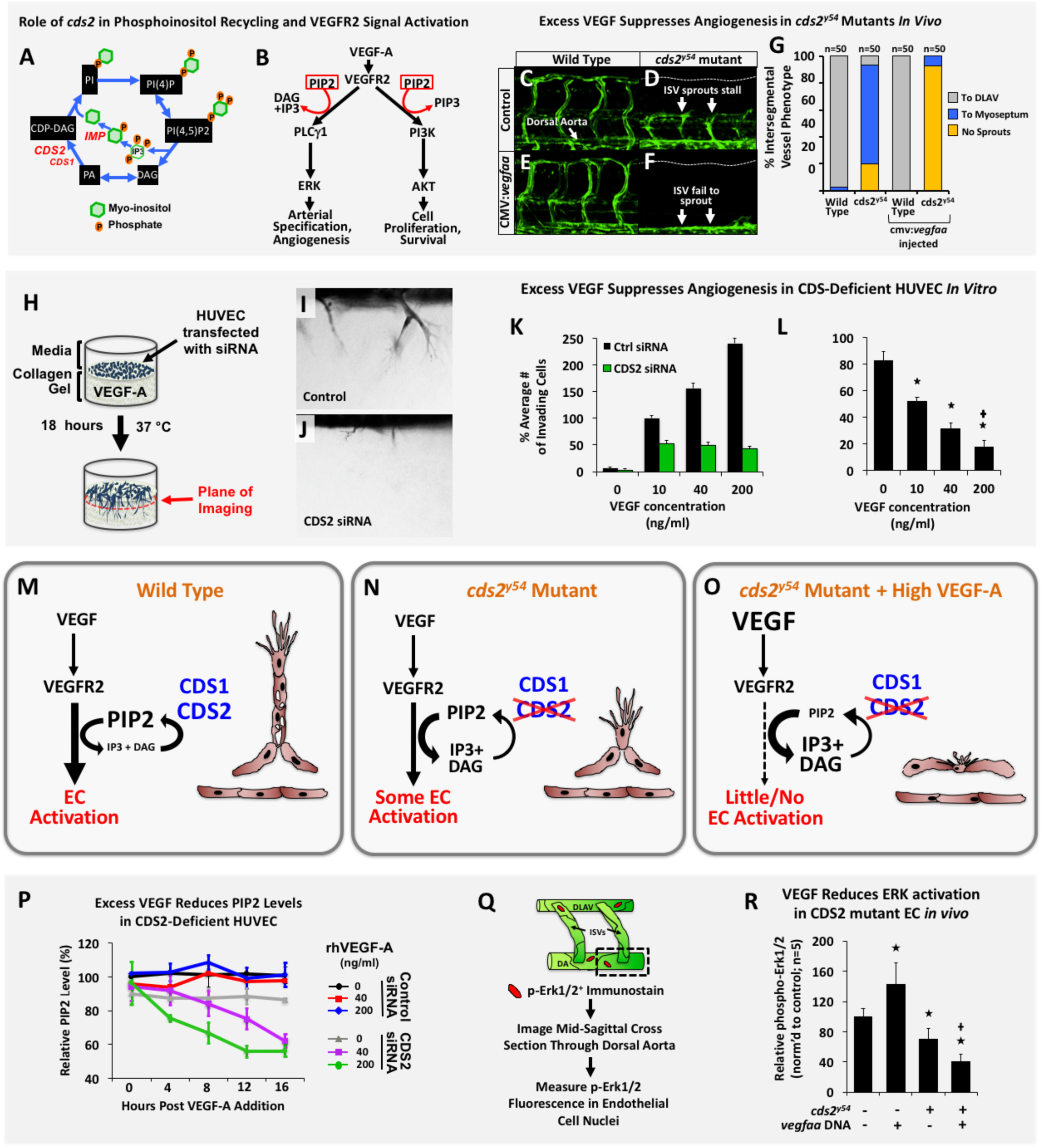
CDS2 dependent angiogenic sprouting defects *in vitro* and *in vivo* are exacerbated by exogenous VEGF-A addition. **(A)** Schematic diagram of phosphoinositide recycling. CDS1, CDS2, and IMP enzymes facilitate regeneration of phosphoinositol after PLCg-dependent consumption of PIP2 (see **Fig. S10** for details). **(B)** VEGFR2 signaling schematic (modified from (*26, 33*)). **(C-G)** Confocal images (C-F) and quantitation (G) of trunk intersegmental vessels (ISV) in 32hpf *Tg(fli-egfp)*^*y1*^ WT siblings (C,E) or *cds2*^*y54*^ mutants (D,F) injected with control (C,D) or CMV:*vegfaa* (E,F) DNA. Bars in G measure ISV that have not sprouted (red), grown halfway up the trunk (blue), or formed a complete ISV (green). **(H-J)** HUVEC 3D invasion assay used to model angiogenesis *in vitro* (H), with representative images from control and CDS2 siRNA treated cultures (I,J). **(K)** Quantification of HUVEC cellular invasion into collagen gels at rhVEGF-A doses indicated. **(L)** Quantification of CDS2 siRNA HUVEC cellular invasion normalized to rhVEGF-A dose-matched controls (see methods); *Significance from control; **+**Significance from individual VEGF-A doses. **(M-O) Phosphoinositide recycling, CDS2, and angiogenesis.** (M) Under normal conditions phosphoinositide recycling maintains PIP2 levels and VEGF signal transduction. (N) Endothelium defective for CDS2 has a reduced capacity to recycle phosphoinositides, but under conditions of initial or low-level VEGF stimulation sufficient PIP2 is regenerated to maintain VEGF signal transduction. (O) During sustained and/or high-level VEGF stimulation, however, PIP2 levels cannot be maintained, leading to a collapse of VEGF signal transduction. **(P)** ELISA quantitation of PIP2 levels in control or CDS2 siRNA treated HUVECs incubated with 0, 40, or 200 ng/ml rhVEGF-A over a 16 hour time course, normalized to levels in initial control siRNA-treated HUVEC without added rhVEGF-A. **(Q)** Diagram illustrating procedure for measurement of phospho-Erk1/2 in trunk endothelial nuclei by immunofluorescence. **(R)** Quantitation of trunk endothelial phospho-Erk1/2 in 30 hpf *cds2*^*y54*^ mutants and WT siblings +/- CMV:*vegfaa* DNA. *Significance from control; **+**Significance from *cds2*^*y54*^ mutant - CMV:*vegfaa* DNA condition.

To further explore the role of phosphoinositide recycling in VEGF signaling, we injected a *Tg(cmv:vegfaa)* transgene (*25*) driving ubiquitous expression of VEGF-A into *cds2*^*y54*^ mutant zebrafish. Instead of promoting increased vessel growth, VEGF-A overexpression in *cds2*^*y54*^ null mutants or *cds2* morpholino (MO) treated embryos results in dramatically reduced angiogenesis-including in the trunk intersegmental vessels, central arteries in the brain, and sub-intestinal vascular plexus (**Fig. 1C-G, Fig. S1**). Longitudinal imaging beginning at 24 hpf reveals that vessels fail to sprout and extend properly (**Fig. S2**). Sensitivity to VEGF is highly specific to *cds2* mutants, as other zebrafish vascular mutants including *plcg1*^*y10*^*, flk1*^*y17*^, and *etsrp*^*y11*^ do not show reduced angiogenesis when injected with *Tg(cmv:vegfaa)* transgene (**Fig. S3**).

Human ECs *in vitro* also exhibit this seemingly paradoxical sensitivity to VEGF stimulation when CDS2 activity is reduced. Two independent siRNA targets were validated in Human Umbilical Vein Endothelial Cells (HUVECs), confirming that both suppressed CDS2 protein levels, disrupted HUVEC invasive capacity *in vitro*, and suppressed p-Erk1/2 and p-Akt signaling downstream of VEGF-A stimulation (**Fig. S4A-C**). CDS2 siRNA treatment reduces HUVEC proliferative capacity but does not induce apoptosis (**Fig. S4D,E**). Exposing CDS2-deficient HUVECs to increasing levels of VEGF stimulation results in decreased migratory capacity in 3-D endothelial invasion assays compared to controls (**Fig. 1H-L**), similar to the reduced angiogenesis noted upon VEGF stimulation in CDS2-deficient zebrafish.

Defects in CDS2 reduce the capacity of ECs to regenerate PIP2 for VEGF signal transduction (**Fig. 1A,B,M,N**) (*9, 10, 12, 16-18, 21, 26, 27*). We hypothesized that while initial and/or low-level VEGF-A stimulation of CDS2-deficient ECs might not substantially affect PIP2 levels (**Fig. 1M,N**), while higher levels and/or sustained VEGF-A stimulation would gradually deplete PIP2 levels, halting VEGF signal transduction (**Fig. 1O**). To examine this directly we measured endogenous PIP2 levels over time in control and CDS2-deficient HUVECs *in vitro* challenged with VEGF-A (**Fig. 1P**). VEGF-A stimulation does not alter endogenous PIP2 levels over a 16-hour time course in control siRNA-treated HUVECs. PIP2 levels are also not significantly reduced over 16 hours in CDS2 siRNA-treated HUVECs in the absence of exogenously added VEGF-A, although the baseline level of PIP2 is lower than in controls. In contrast, however, exogenous VEGF stimulation of CDS2 siRNA-treated HUVECs results in a time-dependent decrease in PIP2 levels, with more rapid reduction in PIP2 observed at higher VEGF-A concentrations.

If PIP2 is similarly reduced in *cds2*-deficient zebrafish we predicted that exogenously supplied PIP2 substrate would rescue the aberrant vascular phenotypes in ECs *in vivo*. Indeed, delivery of a single ‘bolus’ of PIP2 substrate supplied by intravascular injection of PIP2-loaded liposomes largely reverses the angiogenic defects in 48 hpf *cds2*^*y54*^ mutant zebrafish embryos (**Fig. S5**) (*9*).

VEGF signaling downstream from VEGFR2 and PLCg1 is transduced via activation (phosphorylation) of Erk1/2 (**Fig. 1B**). Injection of a *Tg(cmv:vegfaa)* transgene into control zebrafish embryos results in a modest increase in phospho-Erk1/2 levels as assessed by western blot analysis of excised zebrafish trunk tissue (**Fig. S6**), or a more marked increase by immunostaining analysis examining phospho-ERK1/2 specifically in trunk ECs (**Fig. 1Q,R**). CDS-deficient animals show somewhat decreased phospho-Erk1/2, but injection of *Tg(cmv:vegfaa)* transgene into CDS2-deficient zebrafish results in a more dramatic decrease in phospho-Erk1/2, both by western blot (**Fig. S6**) and by immunostaining analysis (**Fig. 1R**). Strikingly, the reduced phospho-Erk1/2 levels in VEGF-overexpressing CDS2-deficient zebrafish are comparable to those in animals injected with *vegfaa* morpholino (**Fig. S6**).

Based on our findings in zebrafish and HUVECs, we hypothesized that phosphoinositide recycling might provide an effective target for anti-angiogenic anti-tumor treatment, as the increased VEGF-A secreted by tumors responding to therapy-induced vascular insufficiency and hypoxia might facilitate anti-angiogenic effects, rather than help to overcome them (**Fig. 1O, Fig. 4**). We used two different murine tumor allograft models, Lewis Lung Carcinoma (LLC) (*23, 28*) and B16-F10 melanoma (*23*), to determine whether systemic or endothelial cell specific suppression of phosphoinositide recycling could specifically inhibit tumor growth and tumor angiogenesis. We employed five different experimental paradigms to target phosphoinositide recycling in these tumor models: direct targeting of CDS2 using separate translation or splice blocking vivoMorpholinos (vMO) (*29*), inhibition of inositol monophosphatase (IMP) activity by the potent chemical inhibitor L-690,488 (*30*), inhibition of IMP activity by lithium chloride (LiCl) treatment, and inducible endothelial-specific genetic deletion of Cds2 via the Cre/Lox system.

We administered murine CDS2 translation-blocking (vMO #1) or splice-blocking (vMO #2) *vivo* Morpholinos (vMO) (*29*), versus a control vMO, into mice allografted with LLC (**Fig. 2A**) or B16 (**Fig. S7A**) tumors. The vMOs were introduced directly into the circulation via tail vein or retro-orbital injection to ensure their bioavailability, beginning two days prior to the implantation of tumor cells into each flank and continuing daily throughout the time course of tumor growth (12-18 days). LLC (**Fig. 2B,C**) or B16-F10 (**Fig. S7B-D**) allografted mice treated with either of the two CDS2 vMOs show decreased tumor volume and decreased final tumor weight compared to control vMO-injected mice. Exposure of isolated B16 or LLC cells to comparable doses of CDS2 vMOs *in vitro* does not alter the proliferation of these cells, suggesting this is not a direct effect on the tumor cells (**Fig. S7E and S8A**). Quantitation of tumor vessel density in CD31 immunostained sections revealed approximately 60-80% reduction in vessel area in either LLC (**Fig. 2D**) or B16 (**Fig. S7F-I**) tumor allografts from CDS2 vMO treated animals compared to controls. Treatment with the vMOs does not change the overall mass of the mice (**Fig. S8B**), and liver tissue from CDS2 vMO treated mice showed no increase in Caspase 3-positive apoptotic cells or decrease in vascularization compared to livers from control vMO treated animals (**Fig. S8C-E**). To further address the specificity of the vMO treatment, we carried out “rescue” experiments using daily intravascular administration of PIP2 liposomes in conjunction with the vMO injections in LLC tumor allograft assays (**Fig. 2E,F**; following a similar logic to that applied to the zebrafish studies in **Fig. S5**). Daily co-injection of PIP2 liposomes into LLC allografted mice treated with the CDS2 vMO reverses the tumor growth defects noted with CDS2 vMO treatment alone (**Fig. 2E,F; Fig. S8F,G**).

**Figure 2.**
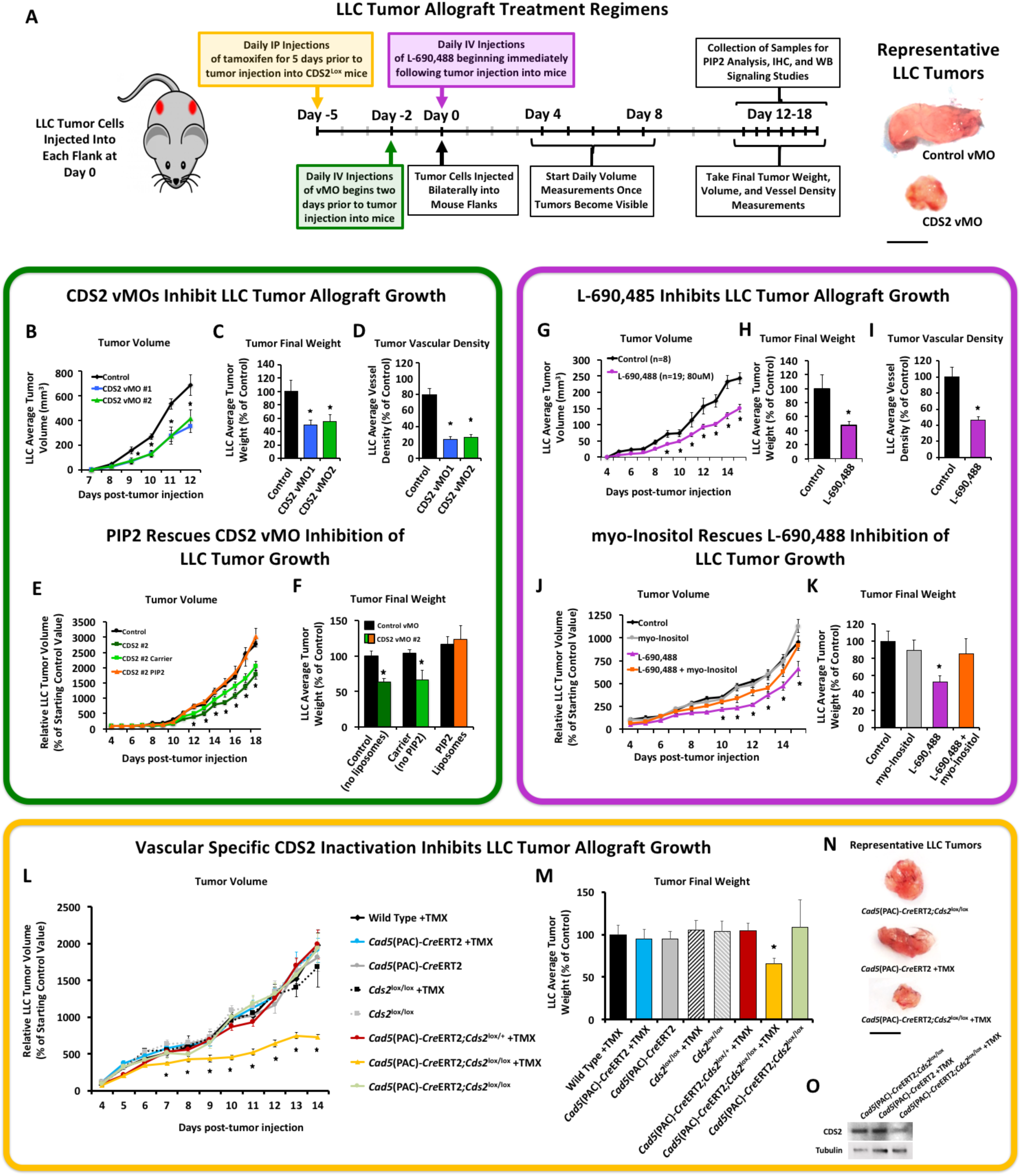
Anti-PIP2 recycling therapies suppress tumor growth. **(A)** Schematic of Lewis Lung Carcinoma (LLC) tumor allograft assay with representative tumor images. Bar = 1 cm. (**B-F**) CDS2 vMO treatment. (**G-K**) L-690,488 small molecule inhibitor treatment. (**L-O**) Endothelial-specific genetic deletion of CDS2. Quantitation of average tumor volume (**B,G,L**), final tumor weight (**C,H,M**) and final tumor vascular density (**D,I**) at 12-15 days post-tumor implantation, in control versus CDS2 vMO or L-690,488 treated animals (B-D vs. G-I) or control or endothelial-specific CDS2 “knockout” animals (L-M). (**E,F**) **PIP2 “rescues” CDS2 vMO tumor inhibition.** Quantitation of LLC average tumor volume (**E**) and final tumor weight (**F**) at 18 days post-tumor implantation in control vMO, CDS2 #2 vMO (no liposomes), CDS2 #2 vMO + carrier liposome (no PIP2), or CDS2 #2 vMO + PIP2 loaded liposome-treated, LLC-allografted mice. Data in (E,F) are normalized to the average starting size of each individual tumor group at day 4, and shown as a percentage of the starting day 4 control (the PIP2 liposome injection start date). (**J,K**) **Myo-inositol “rescues” L-690,488 tumor inhibition.** Quantitation of LLC average tumor volume (**J**) and final tumor weight (**K**) at 15 days post-tumor implantation in control (untreated), myo-inositol-, L-690,488-, or L-690,488 + myo-inositol-treated LLC-allografted mice. Data in (J,K) are normalized to the control condition. (**L-O**) **Endothelial-specific genetic deletion of CDS2 promotes tumor growth inhibition.** Quantitation of LLC average tumor volume (**L**) and final tumor weight (**M**) at 14 days post-tumor implantation in the eight genetic conditions tested. Data in (L) are normalized to the starting day 4 tumor volume of Wild Type + TMX controls. (**N**) Representative tumor images of Vehicle Control (*Cad5*(PAC)- *Cre*ERT2*;Cds2*^lox/lox^); Cre, TMX Control (*Cad5*(PAC)-*Cre*ERT2 +TMX); and endothelial specific Cds2 genetic deletion mice (*Cad5*(PAC)-*Cre*ERT2*;Cds2*^lox/lox^+TMX). (**O**) Representative western blot images of EC protein isolated from tumors for each indicated genetic condition, probed for CDS2 and tubulin as a loading control. Blots demonstrate suppression of the Cds2 protein following TMX treatment in ECs isolated from tumors grown in *Cad5*(PAC)-*Cre*ERT2*;Cds2*^lox/lox^ mice. p ≤ 0.05, ± SEM. *Significance from control. All tumor measurements are ≥ 8 tumors per treatment group. Naming key: Wild type + TMX (no Cre, no Cds2 lox cassette); *Cad5*(PAC)-*Cre*ERT2 +TMX (Cre^iECΔ^ only, +TMX); *Cad5*(PAC)-*Cre*ERT2 (Cre^iECΔ^ only, no TMX vehicle control); *Cds2*^lox/lox^ +TMX (only homozygous Cds2 lox cassette, +TMX, no Cre); *Cds2*^lox/lox^ (only homozygous Cds2 lox cassette, no TMX vehicle control); *Cad5*(PAC)- *Cre*ERT2*;Cds2*^lox/+^ +TMX (Cre^iECΔ^, Cds2 heterozygous lox cassette, +TMX); *Cad5*(PAC)-*Cre*ERT2*;Cds2*^lox/lox^ +TMX (homozygous lox cassette, Cre^iECΔ^, +TMX-experimental deletion group); *Cad5*(PAC)-*Cre*ERT2*;Cds2*^lox/lox^ (homozygous lox cassette, Cre^iECΔ^, no TMX vehicle control)

A similar experimental paradigm was carried out using the potent inositol monophosphatase (IMP) small molecule inhibitor L-690,488 (*30*) in mice with LLC allografted tumors. IMP catalyzes removal of the phosphate group from inositol-1-phosphate to generate myo-inositol, which is then combined with CDP-DAG (the product of the reaction catalyzed by CDS) to regenerate phosphatidylinositol (**Fig. 1A**). LLC tumors from mice treated with L-690,488 show decreased tumor volume and final tumor weight compared to control DMSO-injected mice (**Fig. 2G,H**). Examination of tumor vessel density by CD31 immunostaining of sections revealed approximately 50-60% reduction in vessel area compared to controls (**Fig. 2I**). Treatment with L-690,488 does not affect the overall mass of the mice nor does it decrease normal vessel density in the liver (**Fig. S9**). We carried out “rescue” experiments by co-injecting myo-inositol, the downstream product of the reaction catalyzed by IMP (*9, 24, 30*), along with injections of L-690,488 (**Fig. 2J,K**). Myo-inositol administration effectively reverses tumor growth inhibition caused by L-690,488, suggesting the effects of this drug on tumor growth are specific to inhibition of IMP activity (**Fig. 2J,K**). To further validate our L-690,488 results, we also targeted IMP activity using lithium chloride (LiCl) treatment. Lithium is an effective inhibitor of IMP, although it also has activity against other enzymes and pathways including WNT signaling. We showed previously that LiCl inhibits angiogenesis in developing zebrafish *in vivo* and in EC invasion assays using HUVEC *in vitro*, with the specificity of these effects for IMP verified by myoinositol rescue ^9^. As with L-690,488, we find that treatment with LiCl inhibits tumor growth and tumor angiogenesis in both LLC and B16 allografts, and that these effects are largely reversed by co-administration of myo-inositol (**Fig. S10**).

All of the treatments described above involve systemic inhibition of PI recycling. In order to show that the effects on tumor growth are due to reduced PI recycling in the endothelium, we generated inducible EC-specific Cds2 knockout mice using the *Cad5*(PAC)-*Cre*ERT2 (*31*) EC specific Cre line crossed to Cds2^tm^1^a(KOMP)Wtsi^ (i.e. Cds2^lox/lox^) mice (*32*). Tamoxifen was supplied to the indicated mouse groups for 5 consecutive days prior to the implantation of LLC tumors into the flanks and then suspended for the remainder of the study. LLC tumors from the *Cad5*(PAC)- *Cre*ERT2*;Cds2*^lox/lox^ +TMX (homozygous lox cassette, Cre^iECΔ^, +TMX-experimental deletion group) show decreased tumor volume and decreased final tumor weight compared to all other control groups (**Fig. 2L-N**). Western blot data of EC’s isolated from tumors shows significant suppression of CDS2 protein at the end of the 14 day experimental time course in LLC tumors from *Cad5*(PAC)-*Cre*ERT2*;Cds2*^lox/lox^ +TMX mice (**Fig. 2O**).

The results described above show that anti-phosphoinositide (PI) recycling treatments effectively reduce tumor growth and tumor angiogenesis in LLC or B16 allografted mice without significant effects on the endogenous vasculature of the liver in the same animals. We hypothesized that the distinct effects of anti-PI recycling treatments on tumor versus normal vessels might reflect increased signaling and increased PIP2 substrate utilization in “activated” tumor-associated ECs compared to “quiescent” ECs in endogenous tissues and organs (**Fig. 3A**). We examined this by ELISA quantitation of PIP2 levels in endothelial cells obtained from tumors and lungs of tumor-allografted animals, and by measuring Erk and Akt activation by quantitation of EC fluorescence in immunostained histological sections from tumors and livers of the same mice (**Fig. 3B**). Tumor ECs show substantially higher phospho-Erk and phospho-Akt levels (**Fig. 3C**), as well as higher PIP2 levels (**Fig. 3D**) than liver or lung ECs (respectively). The increased angiogenic signaling in tumor-associated ECs supports the idea that systemic administration of appropriate doses of PI recycling inhibitors can have specific effects on “activated” tumor ECs but not on normal “quiescent” ECs.

**Figure 3.**
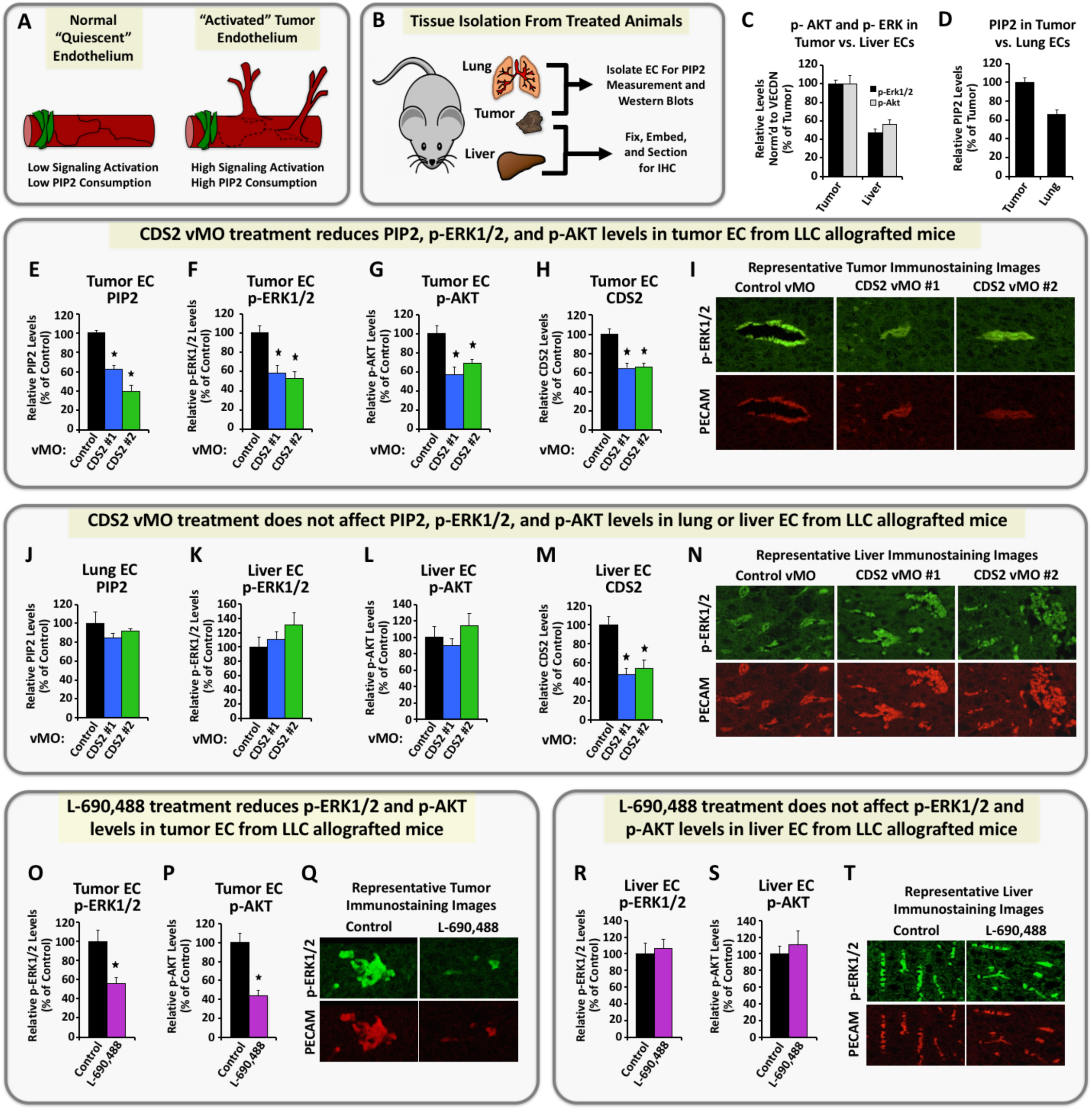
Systemic anti-PIP2 recycling therapies suppress VEGFR2 downstream signaling in LLC tumor models. **(A)** Model for signaling in activated versus quiescent endothelium. **(B)** Diagram illustrating tissues isolated from mice and their use. **(C)** Quantification of phospho-Erk1/2 (black bars) and phospho-Akt (gray bars) protein levels in immunostained liver or tumor tissue from LLC allografted control animals. **(D)** ELISA measurement of PIP2 levels in lung or tumor endothelial cells from LLC allografted control animals. **(E-N) CDS2 vMO-treated, LLC-allografted mice.** Quantification of PIP2 levels by ELISA in tumor (**E**) or lung (**J**) endothelial cells isolated from the same LLC allografted animals. Quantification of phospho-Erk1/2 (**F,K**), phospho-Akt (**G,L**) and CDS2 (**H,M**) protein levels from immunostained sections of tumor (**F-H**) or liver (**K-M**) tissue collected from the same LLC allografted animals. **(I,N)** Representative images of tumor (I) or liver (N) sections from control or CDS2 vMO treated mice, double-immunostained with phospho-Erk1/2 antibody (green images) and PECAM antibody (red images). **(O-T) L-690,488-treated, LLC-allografted mice.** Quantification of phospho-Erk1/2 (**O,R**) and phospho-Akt (**P,S**) protein levels from immunostained sections of tumor (**O,P**) or liver (**R,S**) tissue collected from the same LLC allografted animals. **(Q,T)** Representative images of tumor (Q) or liver (T) sections from control (DMSO) or L-690,488 treated mice, double-immunostained with phospho-Erk1/2 antibody (green images) and PECAM antibody (red images). p ≤ 0.05, ± SEM. *Significance from control.

To determine whether systemic inhibition of PI recycling indeed preferentially reduced signaling in tumor-associated ECs as compared to quiescent endogenous ECs, we analyzed ECs from tumors (**Fig. 3E-I,O-Q**) or livers/lungs (**Fig. 3J-N,R-T**) of the same CDS2 vMO (**Fig. 3E-N**) or L-690,488 (**Fig. 3O-T**) treated, LLC-allografted mice. ECs from LLC tumors have markedly reduced PIP2, p-Akt, p-Erk1/2, and CDS2 protein levels following CDS2 vMO treatment (**Fig. 3E-I; Fig. S11G-L**). In contrast, EC from lungs and livers of the same CDS2 vMO treated mice show no reduction in PIP2, p-Akt, or p-Erk1/2 levels (**Fig. 3J-L,N; Fig. S11B-E**), despite having comparable reduction in CDS2 protein levels (**Fig. 3M; Fig. S11A,F**). Similar differential effects on tumor versus liver EC p-Erk1/2 and p-Akt activation were noted in mice treated with the IMP inhibitor L-690,488 (**Fig. 3O-T**). Taken together, these data show that systemic inhibition of PI recycling depletes PIP2 levels and causes angiogenic defects in the tumor-associated vasculature, but has little effect on the pre-existing stable lung or liver vasculature of the same animals.

Our findings highlight the important role that phosphoinositide recycling plays in tumor angiogenesis, and suggest that targeted inhibition of PI recycling may provide a useful new anti-angiogenic modality for the treatment of cancer. As noted above, the effectiveness of anti-angiogenic therapies targeting VEGF ligand, receptor, or downstream signal transduction has been hampered by the ability of tumors to overproduce VEGF and other cytokines, overcoming endothelial inhibition (**Fig. 4A**). The paradoxical VEGF sensitivity of ECs subjected to anti-PI recycling treatments suggests that high levels of tumor-secreted VEGF may facilitate the effectiveness of anti-PI recycling therapies rather than overcome them (**Fig. 4B**), although further studies will be needed to further validate this exciting possibility.

**Figure 4.**
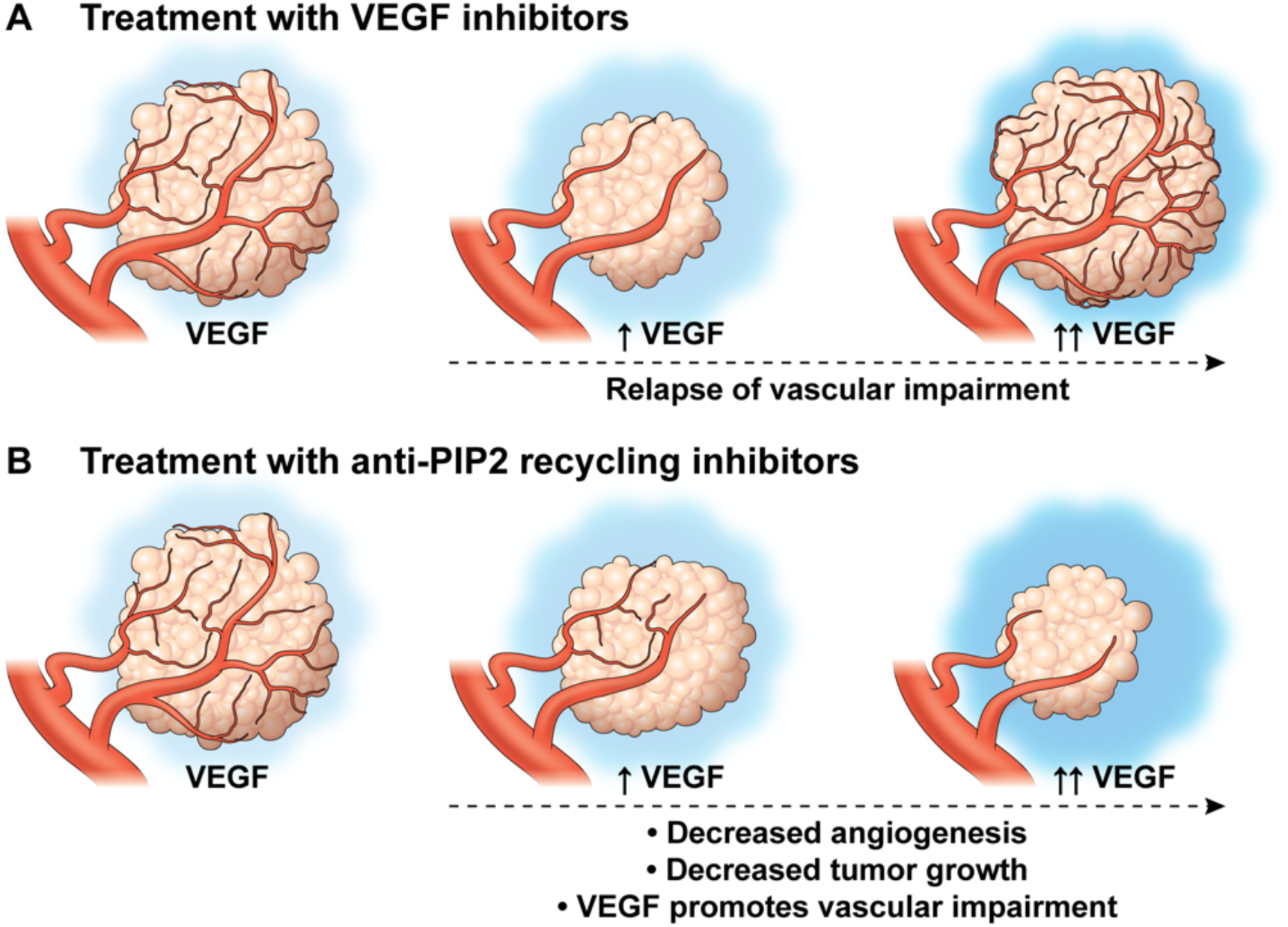
Proposed model for inhibition of tumor growth with anti-VEGF versus anti-PIP2 recycling therapies. Currently available anti-angiogenic therapies targeting VEGF (top) lead to an initial decrease in tumor growth and angiogenesis. However, production of high levels of pro-angiogenic ligands by tumors results in a relapse in vascular impairment and return of tumor growth. Our results indicate that targeting PI recycling leads to reduced tumor vasculature and tumor growth (bottom), and further suggest that the effects on tumor angiogenesis and tumor growth may be strengthened, not overcome, as tumor production of VEGF increases.

## ACKNOWLEDGEMENTS

The authors would like to thank members of the Weinstein laboratory for their critical comments on this manuscript. This work was supported by the intramural program of the National Institute of Child Health and Human Development, National Institutes of Health (NIH; B.M.W.), the intramural program of the National Institute of Dental and Craniofacial Research, National Institutes of Health (NIH; J.S.G), and K99/R00 Pathway to Independence Award (4R00HL125683 – 02; A.N.S.).

## AUTHORSHIP

Contribution: A.N.S, C.M.M., Z.W., O.M.F, M.F.M, M.C.B, V.N.P., S.A.P., D.C., A.E.D., J.J.Y, and T.M.K. performed experiments; A.N.S, C.M.M., Z.W., O.M.F, M.F.M, S.A.P, V.N.P., S.A.P., D.C., A.E.D., J.J.Y, T.M.K., G.E.D., J.S.G., and B.M.W. analyzed results and made the figures; A.N.S, C.M.M., V.N.P., G.E.D., J.S.G., and B.M.W. designed the research and wrote the paper.

Conflict-of-interest disclosure: The authors declare no competing financial interests. Correspondence: Brant M. Weinstein, Program in Genomics of Differentiation, National Institute of Child Health and Human Development, National Institutes of Health, 6 Center Dr. Bethesda, MD 20892.

## SUPPLEMENTARY MATERIAL

**Supplementary Figure 1.**
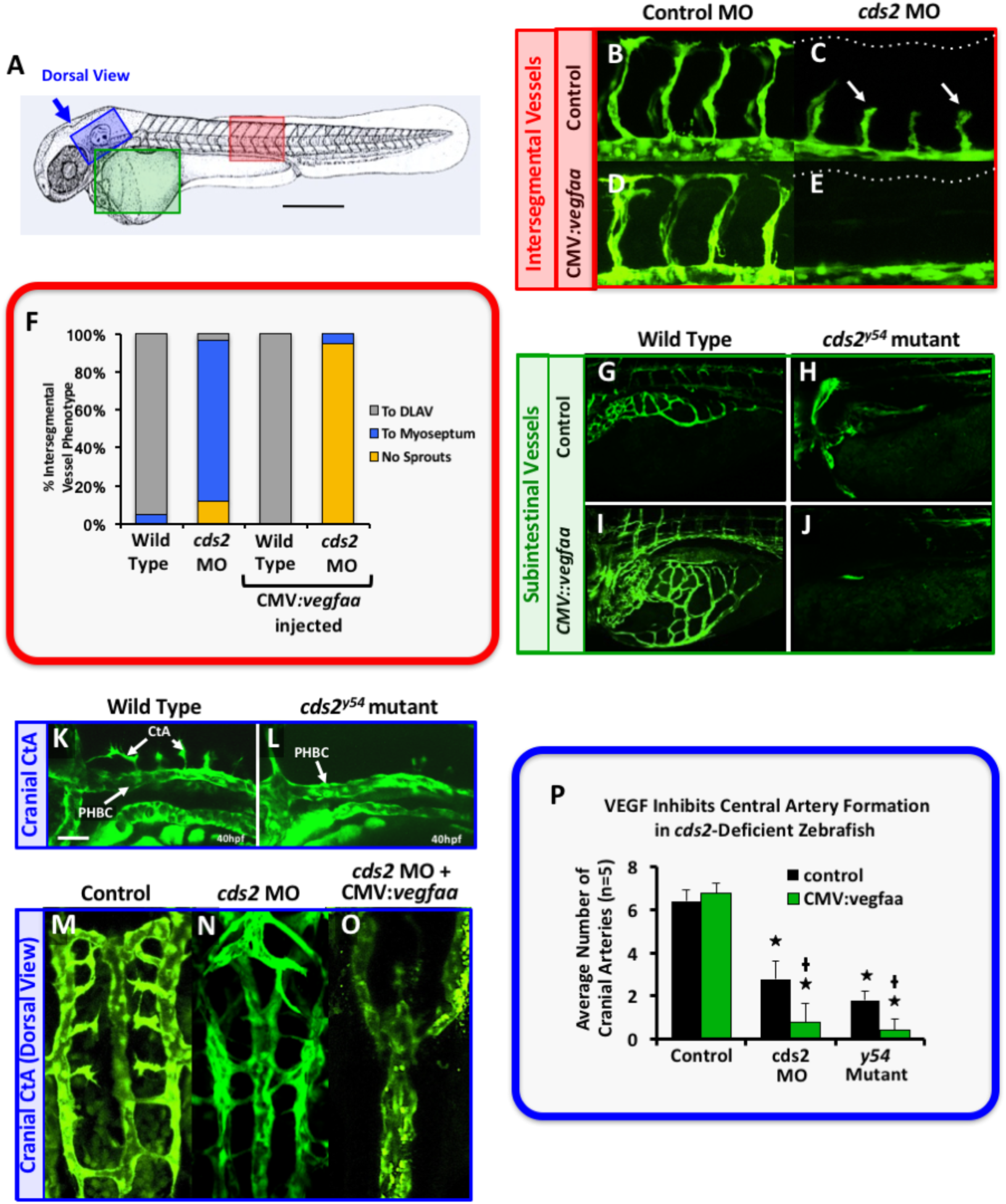
*cds2* mutants and morphants challenged with exogenous *vegfaa* have enhanced angiogenic defects in multiple vascular beds. **(A)** Diagram of a zebrafish embryo with the red box highlighting the region imaged in panels b-f, green box highlighting the region imaged in panels g-j, and blue box highlighting the region imaged and quantified in panels k-p. **(B-E)** Confocal images of trunk ISVs in 32 hpf *Tg(fli-EGFP)*^*y1*^ zebrafish control morpholino (MO, B) or *cds2* MO (C) injected with or control (D) or CMV:*vegfaa* DNA (E). Injection of *cds2* MO elicits a phenotype indistinguishable from *cds2* mutants, including stalling of ISV angiogenesis (white arrows in panel C), and exacerbated inhibition of angiogenesis in the presence of CMV:*vegfaa* DNA (E). **(F)** Quantitation of the ISV phenotypes of 32 hpf *Tg(fli-EGFP)*^*y1*^ control or *cds2* MO animals injected with control or CMV:*vegfaa* DNA. Bars show the percentage of ISV that have failed to sprout (red), ISV that have grown only up to the horizontal myoseptum half-way up the trunk (blue), and ISV that have grown to the dorsal trunk to form the DLAV (green). **(G-J)** Confocal images of 3 dpf *Tg(fli-EGFP)*^*y1*^ subintestinal vessels in wild type (G,I) or *cds2*^*y54*^ mutant (H,J) animals in the absence (G,I) or presence (H,J) of injected CMV:*vegfaa* DNA. **(K-O)** Confocal images of hindbrain vessels in 40 hpf *Tg(fli-EGFP)*^*y1*^ zebrafish. (K,L) Lateral view of the hindbrain vasculature in wild type (K) and *cds2*^*y54*^ mutant (L) animals, showing defects in formation of angiogenic CtA in *cds2*^*y54*^ mutants (*1*). (M-O) Dorsal view of the hindbrain vasculature in animals injected with control MO (m), *cds2* MO (n), or *cds2* MO + CMV:*vegfaa* DNA (o). **(P)** Quantitation of the number of CtA in the hindbrains of 40 hpf *Tg(fli-EGFP)*^*y1*^ wild type control, *cds2* MO injected, or *cds2*^*y54*^ mutant animals that were either un-injected (black bars) or injected with CMV:*vegfaa* DNA (green bars). Overexpressing *vegfaa* similarly exacerbates CtA defects in both *cds2* morpholino-injected and *cds2*^*y54*^ mutant animals. Abbreviations: Posterior hindbrain channel (PHBC), Central Arteries (CtA). Rostral is to the left and dorsal is up in all image panels except panels M-O. In panels M-O, rostral is up and left is to the left. Scale bars = 100 µm. p ≤ 0.05; *Significance from control; +Significance from paired control.

**Supplementary Figure 2.**
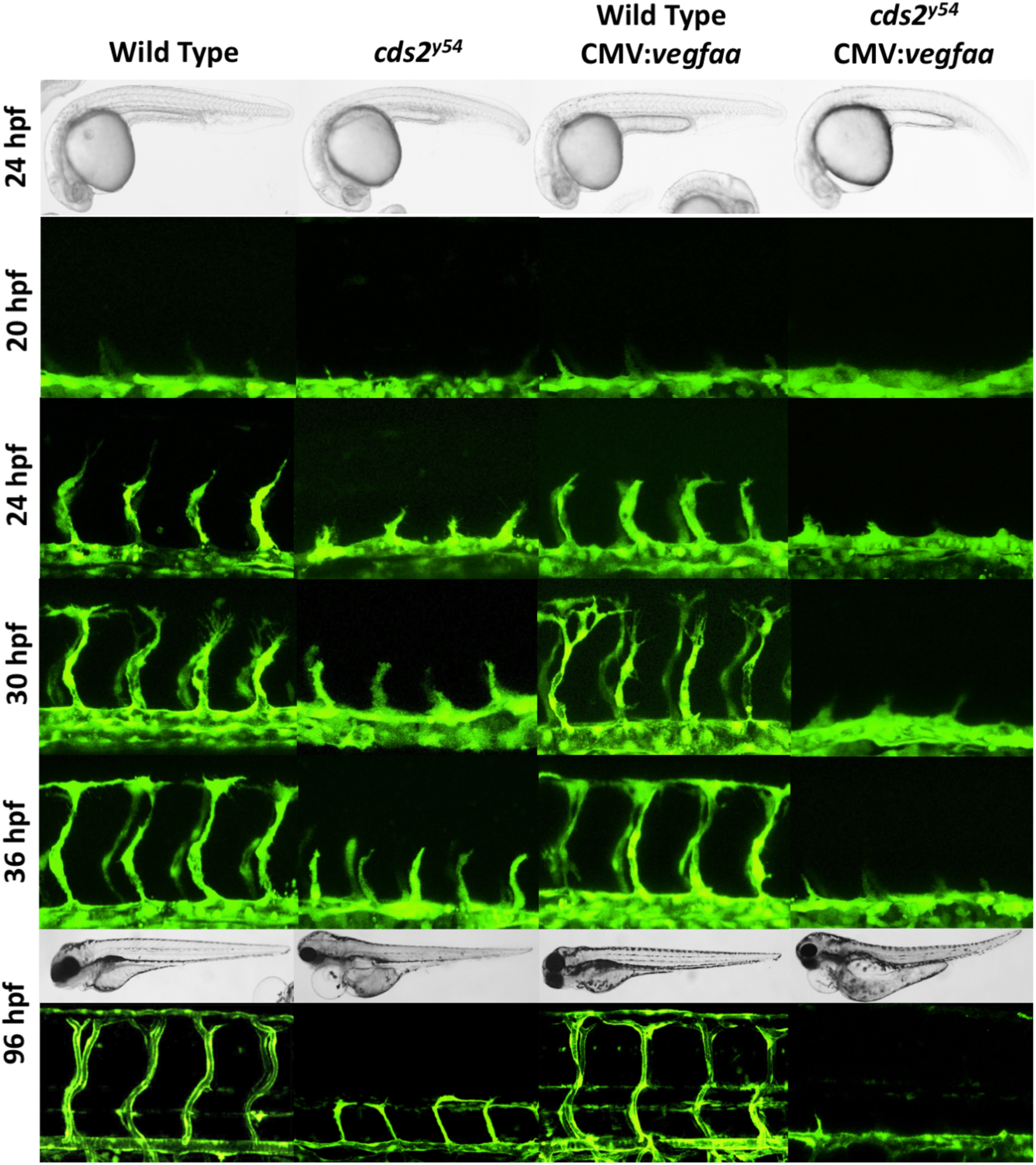
Exogenous *vegfaa* stimulation causes stalling of angiogenic sprouts, not vascular regression. Confocal images of trunk vascular phenotypes in control and CMV:*vegfaa* injected wild type or *cds2*^*y54*^ mutants at 20, 24, 30, 36 and 96 hpf. Column I: control DNA, wild type, Column II: control DNA, *cds2*^*y54*^ mutant, Column III: CMV:*vegfaa* DNA, wild type, Column IV: CMV:*vegfaa* DNA, *cds2*^*y54*^ mutant.

**Supplementary Figure 3.**
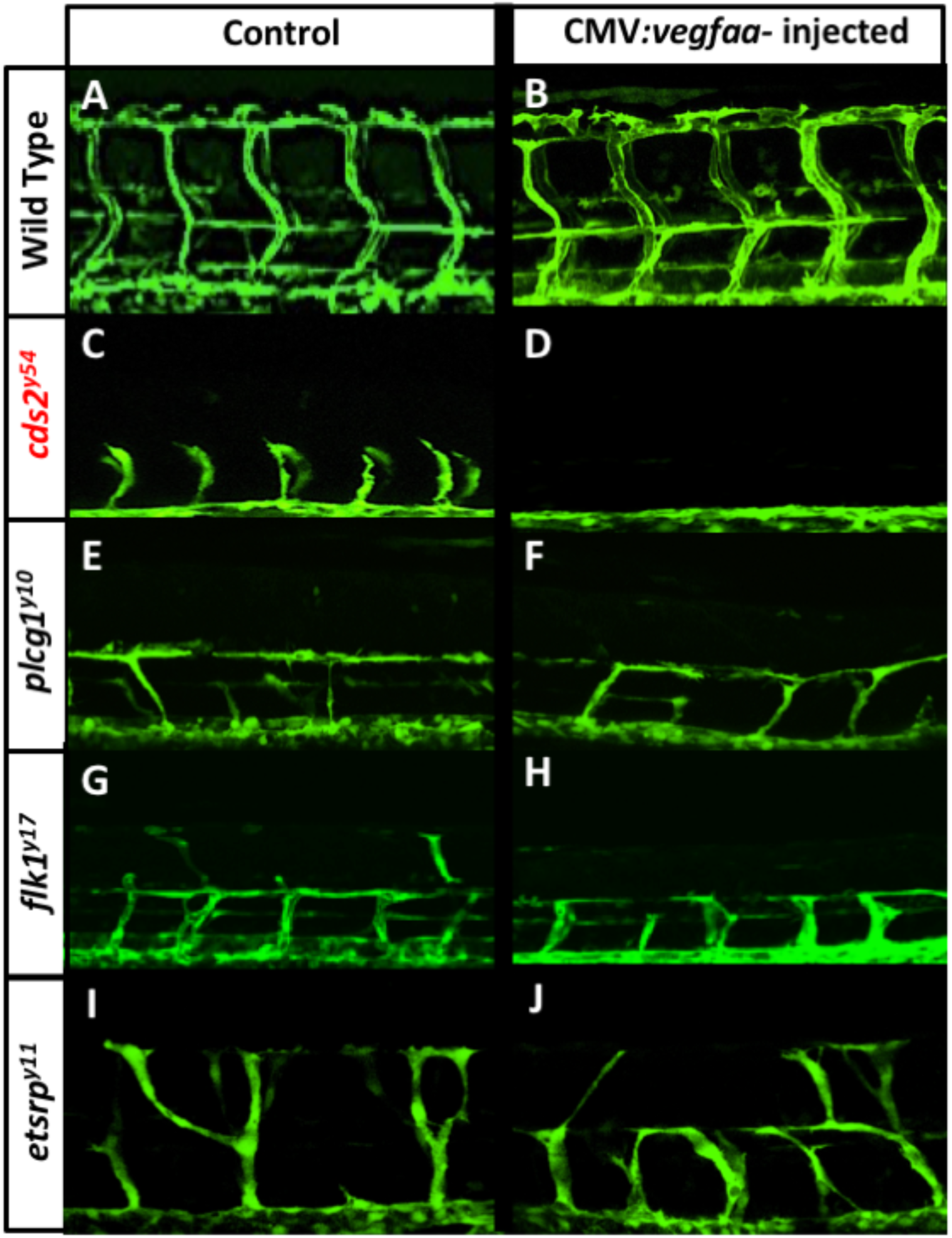
Exogenous *vegfaa* stimulation uniquely exacerbates vascular defects in *cds2*^*y54*^ mutants. **(A-J)** Confocal images of trunk vascular phenotypes in control (A,C,E,G,I) and CMV:*vegfaa* injected (B,D,F,H,J) 36 hpf *Tg(fli-EGFP)*^*y1*^ transgenic zebrafish embryos that were either wild type (A,B), *cds2*^*y54*^ mutant (C,D), *plcγ1*^*y10*^ mutant (E,F), *flk1*^*y17*^ mutant (G,H), or *etsrp*^*y11*^ mutant (I,J). Only *cds2*^*y54*^ mutants show exacerbated vascular defects following injection of CMV:*vegfaa*.

**Supplementary Figure 4.**
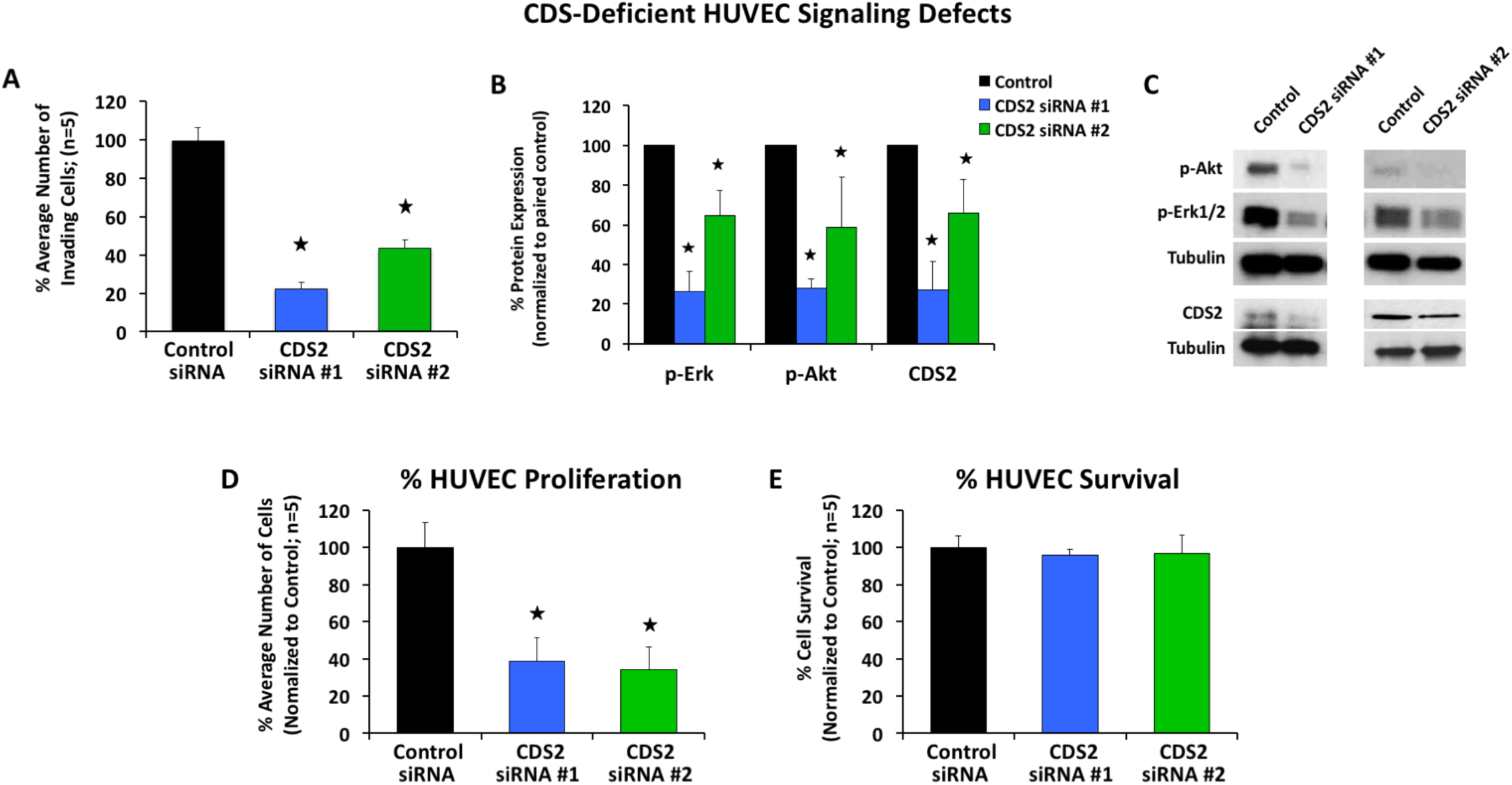
*In vitro* defects of CDS2 deficient endothelium and CDS2 siRNA validation. **(A)** Two single siRNA targets of CDS2 were chosen for analysis. HUVECs treated with control or the CDS2 siRNAs were plated on top of collagen type I gel and cellular invasion into the collagen gel quantified at 18 hours. **(B,C)** Western blot analysis of HUVECs treated with control or the individual CDS2 siRNA targets. Lysates were collected following incubation with rhVEGF-A and p-Erk1/2, p-Akt and CDS2 protein levels analyzed. Quantification (B) and representative blots (C) for each of the siRNAs are shown. Analysis of CDS2 protein levels shows approximately 75-80% protein suppression using CDS2 siRNA target #1, so it was chosen for use in all of the main text figures. **(D,E)** Proliferation (D) and apoptosis (Total cell number – Caspase 3 positive cells; E) were analyzed in cells treated with control or the individual CDS2 siRNA targets. All data is represented as a percentage of the control siRNA treated cells. p ≤ 0.05, ± SD; *Significance from control.

**Supplementary Figure 5.**
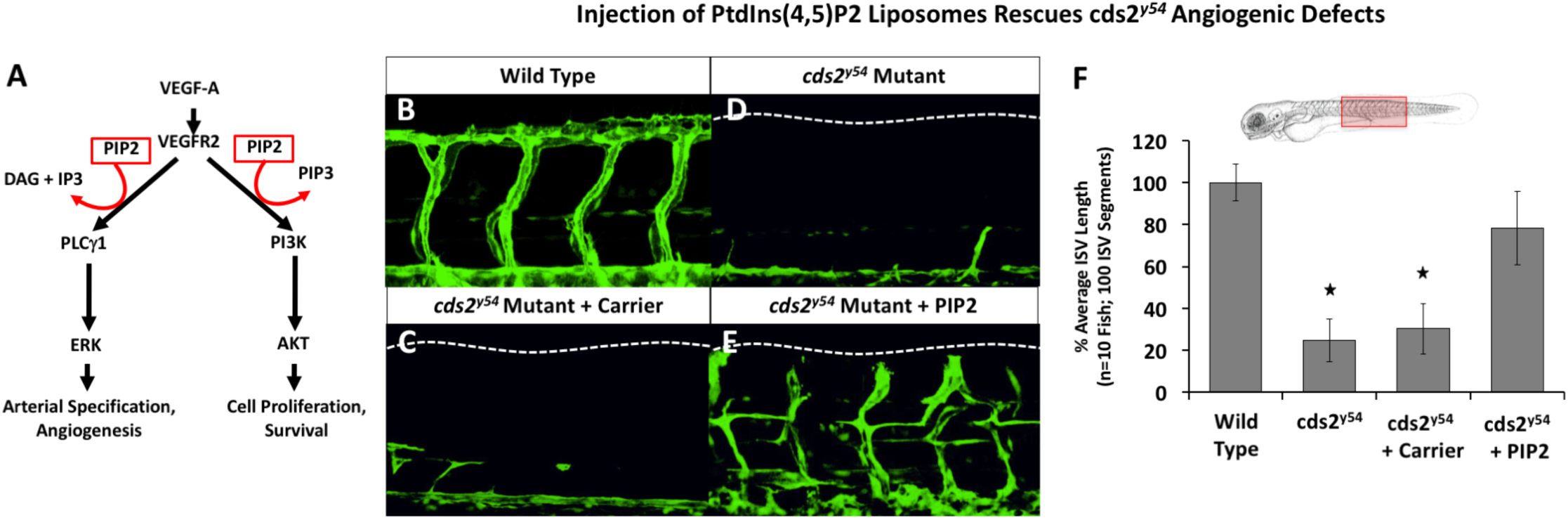
PIP2 liposome injection rescues *cds2*^*y54*^ angiogenic defects. **(A)** Schematic demonstrating the requirement of for VEGF-mediated angiogenesis. **(B-E)** Confocal images of ISV in 72 hpf *Tg(fli-EGFP)*^*y1*^ wild-type animals (B) or *cds2*^*y54*^mutant siblings (C-E) that were uninjected (D) or subjected to a single intravascular injection at 48 hpf with liposome carrier (C) or liposome carrier + PIP2 (E). **(F)** Quantitation of average ISV length in *Tg(fli-EGFP)*^*y1*^ wild-type (column 1) or *cds2*^*y54*^ mutant sibling animals that were either uninjected (column 2) or subjected to a single intravascular injection at 48 hpf with liposome carrier (column 3) or liposome carrier + PIP2 (column 4). Percent average ISV length with wild type average length set to 100%. Schematic indicates the site for analysis within the zebrafish trunk. p ≤ 0.05, ± SEM; *Significance from wild type.

**Supplementary Figure 6.**
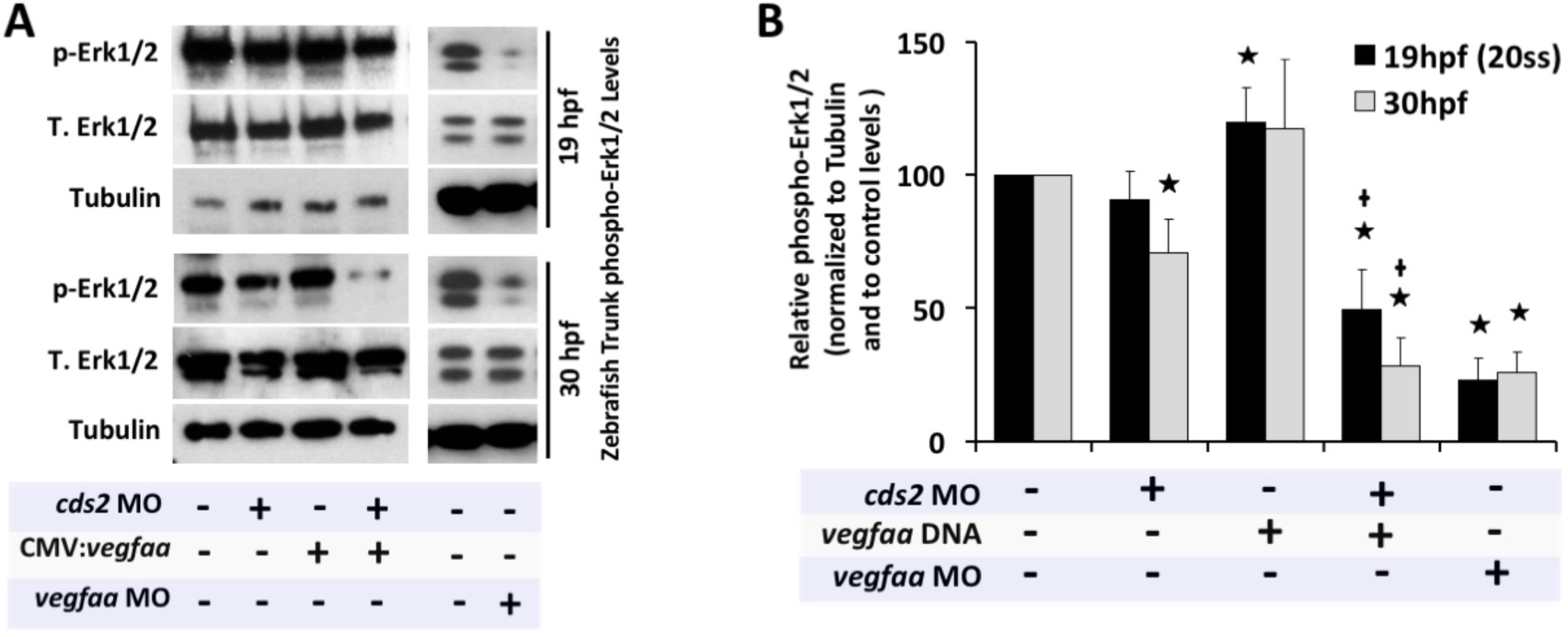
VEGF-A inhibits Erk1/2 signaling in CDS2 deficient endothelium. **(A)** Representative western blots and **(B)** quantitation of Erk1/2 phosphorylation in zebrafish trunk tissue collected at 19 hpf (black bars) or 30 hpf (green bars) from animals injected with (i) control MO, (ii) *cds2* MO, (iii) CMV:*vegfaa* DNA, (iv) *cds2* MO + CMV:*vegfaa* DNA, or (v) *vegfaa* MO. Bars indicate the ratio of pErk1/2 to tubulin levels for each condition, normalized to the phospho-Erk1/2/tubulin levels of the control MO injected animals (set at 100%). p ≤ 0.05, ± SEM; *Significance from control; **+**Significance from *cds2*^*y54*^ mutant or MO alone condition.

**Supplementary Figure 7.**
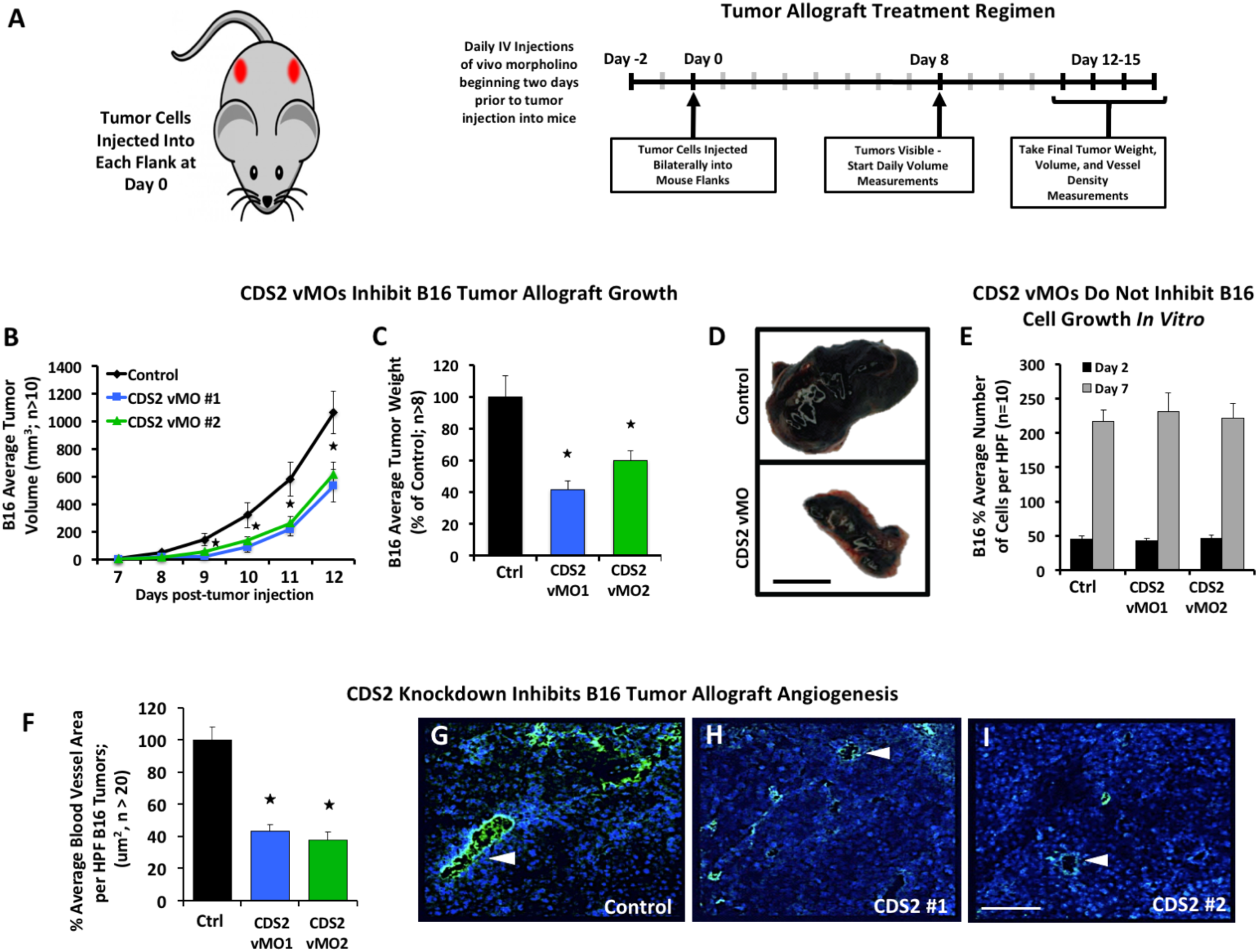
Systemic CDS2 suppression inhibits tumor growth in B16/F10 allograft models. **(A)** Schematic of tumor allograft assay. B6/C57 adult mice received daily intravenous injections of control vivoMorpholino (vMO) or vMO targeting one of two independent sites in the CDS2 gene, starting two days prior to tumor implantation. B16/F10 melanoma tumor cells were injected into each flank of the mice at day 0 and tumors allowed to develop for 12-15 days. Volume measurements were taken daily when tumors became visible with final tumor volumes and weights, mouse weight, and tumor blood vessel density measurements taken at the termination of the experiment. **(B-E)** B16 melanoma tumor volume (B), final tumor weight (C), representative B16 tumors from control versus vMO treated animals (D), and number of B16 melanoma cells present after 2 or 7 days of growth *in vitro* (E). Quantitation of average tumor volume (in mm^3^) of B16 tumors, averaging the measurements of ≥ 10 tumors per treatment group. Quantitation of average tumor weight (in mg) of B16 tumors at 12 days post-tumor implantation, averaging the measurements of ≥ 8 tumors per treatment group. **(E)** Quantitation of the average number of B16 tumor cells per high power field, measured 2 or 7 days (approximately 3 and 14 division cycles) after seeding 5,000 cells per well, with vMO doses comparable to those administered to the mice. p ≤ 0.05, ± SEM. *Significance from control; Bar = 1 cm in panel D. **(F)** Quantitation of B16 tumor vessel density (F) in CDS2 vMO-versus control vMO-treated mice. **(G-I)** Representative images of CD31/PECAM labeled (green) B16 tumor sections from control vMO (G) and CDS2 vMO (H,I) treated mice. White arrowheads indicate sites of CD31/PECAM positive blood vessel labeling. Bar = 100 μm. p ≤ 0.05, ± SEM. *Significance from control.

**Supplementary Figure 8.**
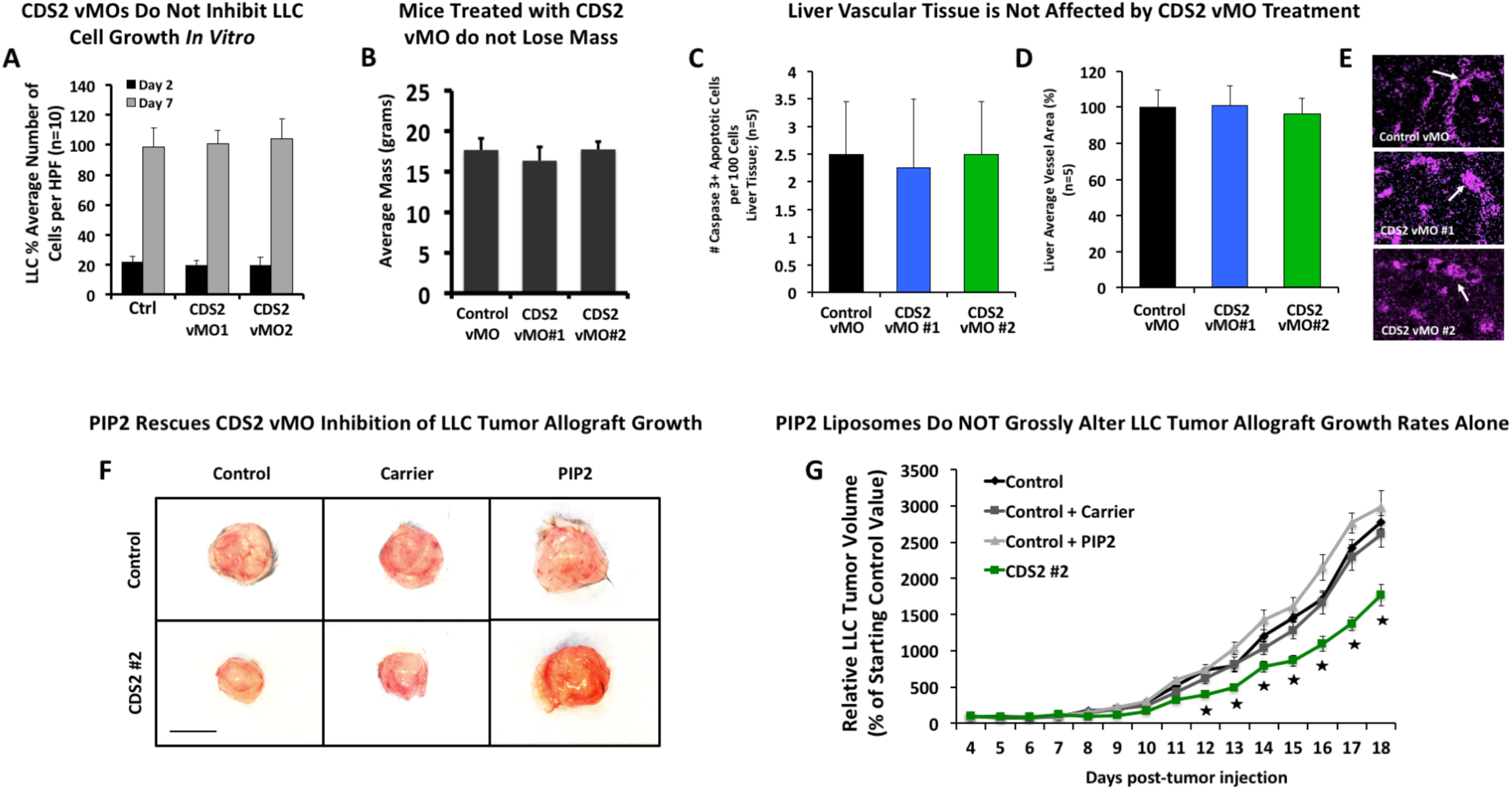
Systemic CDS2 suppression inhibits tumor growth in LLC allograft tumor models but does not affect normal tissue. **(A)** Quantitation of the average number LLC tumor cells per high power field, measured 2 or 7 days (approximately 3 and 14 division cycles) after seeding 5,000 cells per well, with vMO doses comparable to those administered to the mice. **(B)** Quantitation of final mass (in grams) of mice treated with either control or CDS2-specific vMOs. No significant decrease in mass was noted following treatment with CDS2 vMOs. **(C)** Average number of caspase 3+ cells per 100 cells in liver tissue collected from control versus CDS2 vMO treated animals. No significant change was noted. **(D,E)** Analysis of vessel density in liver tissue collected from control versus CDS2 vMO treated animals. (D) Quantification and (E) representative images of CD31/PECAM labeled vessels (purple, highlighted by white arrows) are shown for comparison. Data is represented as a percentage of control. No significant changes were noted. **(F)** Representative images of tumors collected from control vMO treated animals or CDS2 vMO treated animals co-administered nothing, carrier or PIP2 liposomes at 18 days post-tumor implantation. Bar =1 cm. **(G)** Average percent volume measurements (in mm^3^) of control, control + carrier and control + PIP2 versus CDS2 vMO #2 treatment conditions. p ≤ 0.05, ± SEM. *Significance from control.

**Supplementary Figure 9.**
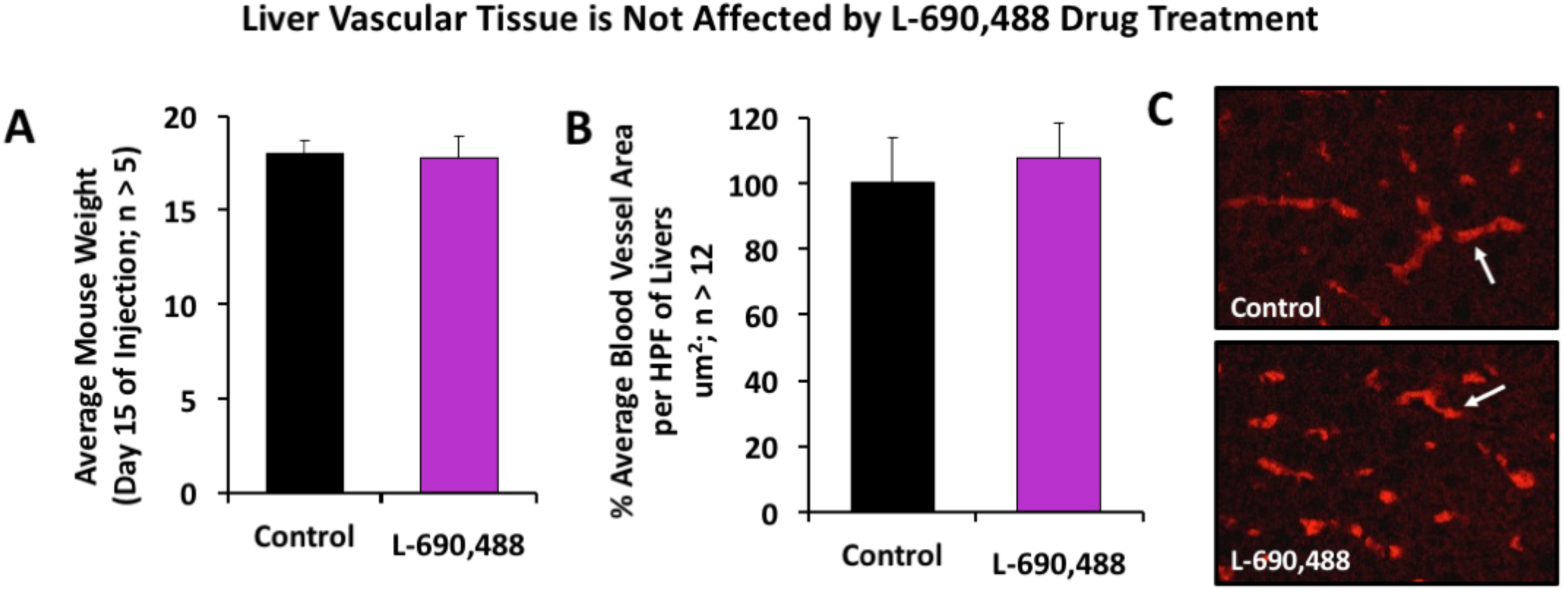
L-690,488 treatment does not affect normal tissue. **(A)** Quantitation of final mass (in grams) of mice treated with either control (DMSO) or L-690,488. No significant decrease in mass was noted following treatment with the chemical inhibitor. **(B,C)** Analysis of vessel density in liver tissue collected from control (DMSO) versus L-690,488 treated animals. (B) Quantification and (C) representative images of CD31/PECAM labeled vessels (red, highlighted by white arrows) are shown for comparison. Data is represented as a percentage of control. No significant changes were noted.

**Supplementary Figure 10.**
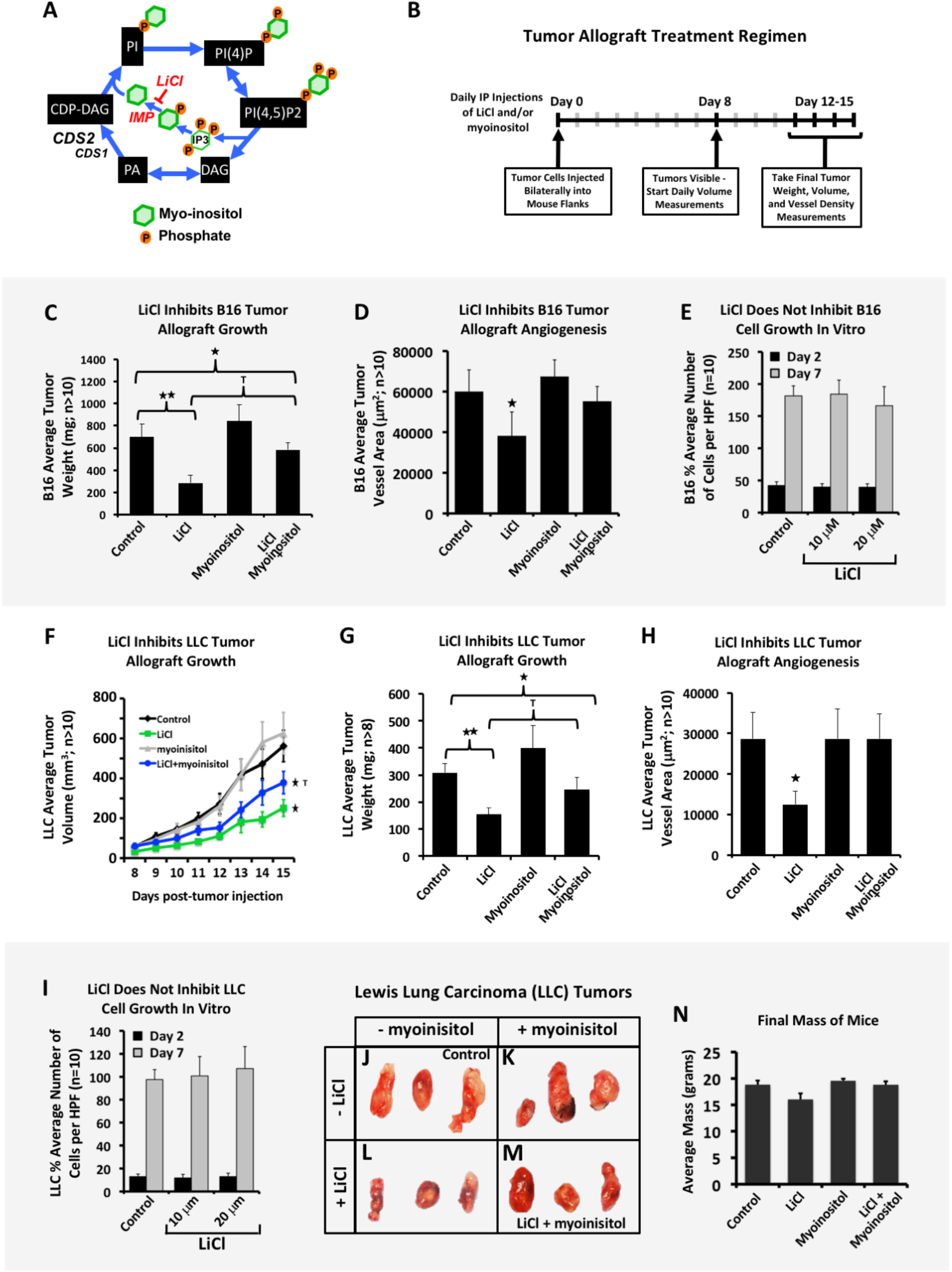
Lithium chloride inhibits B16 and LLC tumor growth and tumor vascularization. **(A)** Schematic diagram of the blocking effect of LiCl on myoinositol-monophosphatase (IMPase), an enzyme necessary for recycling the myoinositol rings required to regenerate PI and its isoforms. Addition of myoinositol, the product of the reaction catalyzed by IMPase, can be used to “rescue” the effects of IMPase suppression, revealing IMPase-specific effects of LiCl as opposed to other, off-target effects (that are not rescued by myoinositol). **(B)** Schematic of tumor allograft assay for LiCl treatment experiments. B16 melanoma or Lewis Lung Carcinoma (LLC) tumor cells were injected into each flank of the mice at day 0 and tumors were allowed to develop for 12-15 days with daily IP injections of LiCl and/or myoinositol. Tumor volume measurements were taken daily starting at approximately day 8, when tumors became visible, with final tumor volumes and weights, mouse weight, and tumor blood vessel density measurements taken at the termination of the experiment. **(C-E)** B16 melanoma final tumor weight (C), tumor vessel density (D), and number of B16 melanoma cells present after 2 or 7 days of growth *in vitro* (E). **(F-I)** LLC tumor volume (F), final tumor weight (G), tumor vessel density (H), and number of LLC cells present after 2 or 7 days of growth *in vitro* (I). **(F)** Quantitation of average LLC tumor volume (in mm^3^), averaging the measurements of ≥10 tumors per treatment group. **(C,G)** Quantitation of average tumor weight (in mg) of B16 (C) or LLC (G) tumors at 12 days post-tumor implantation, averaging the measurements of ≥8 tumors per treatment group. **(D,H)** Quantitation of B16 (D) or LLC (H) tumor vessel density in CDS2 vMO-versus control vMO-treated mice (see methods). **(E,I)** Quantitation of the average number of B16 (E) or LLC (I) tumor cells per high power field, measured 2 or 7 days (approximately 3 and 14 division cycles) after seeding 5,000 cells per well, with a LiCl dose comparable to that administered to the mice (10µM). **(J-M)** Representative images of LLC tumors from control (J), myoinositol treated (K), LiCl treated (L), or LiCl + myoinositol treated (M) mice. **(N)** Final total mass of mice from panel G. Abbreviations: myoinositol-monophosphatase (IMPase), Diacylglycerol (DAG), Phosphoinositol (PI), Phosphatidic acid (PA), CDP-DAG synthetase (CDS). P ≤ 0.05, ±SEM; *Significance from control; ^T^Significance from LiCl treatment condition. **Abbreviations:** myoinositol-monophosphatase (IMPase), Diacylglycerol (DAG), Phosphatidylinositol (PI), Phosphatidic acid (PA), inositol triphosphate (IP3), Phospholipase C-gamma (PLCγ), phosphatidylinositol-4,5-bisphosphate (PIP2), CDP-diacylglycerol synthetase (CDS).

**Supplementary Figure 11.**
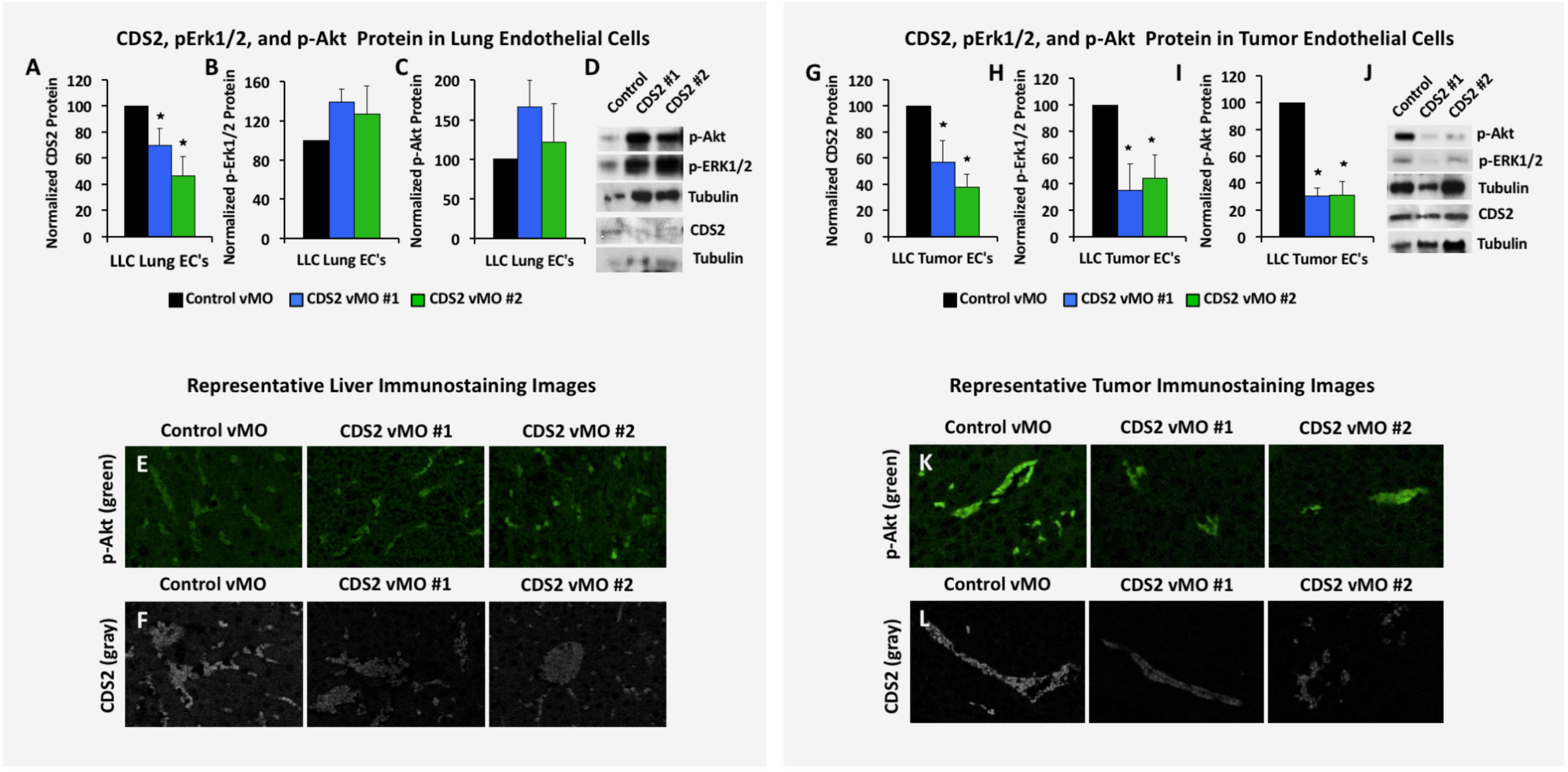
Systemic CDS2 suppression inhibits VEGFR2 downstream signaling in LLC tumor models. **(A-D)** Quantification of CDS2 (A), pErk1/2 (B), and pAkt (C) protein levels in ECs isolated from the lungs of LLC allografted animals. (D) Representative western blot images from lung isolated ECs. CDS2 western blot for lung ECs is shown from a separate run of the same samples and is presented with its respective tubulin loading control. **(E,F)** Representative images of liver cross sections from the same control vMO or CDS2 vMO treated mice, immunostained with pAkt antibody (E, green images) or CDS2 antibody (F, gray images). **(G-J)** Quantification of CDS2 (G), pErk1/2 (H), and pAkt (I) protein levels in ECs isolated from LLC allografted tumors. (J) Representative western blot images from tumor isolated ECs. CDS2 western blot for tumor ECs is shown from a separate run of the same samples and is presented with its respective tubulin loading control. p ≤ 0.05, ±SEM. *Significance from control. **(K,L)** Representative images of LLC tumor cross sections from the same control vMO or CDS2 vMO treated mice, immunostained with pAkt antibody (K, green images) or CDS2 antibody (L, gray images).

## MATERIALS AND METHODS

### Zebrafish Methods

Zebrafish (Danio rerio) embryos were raised and maintained as described (*2, 3*). The *Tg(fli1a:EGFP)*^*y1*^ transgenic zebrafish line was previously described(*4*). The *cds2*^*y25*^ and *cds2*^*y54*^ mutants were identified in an F3 genetic screen in the *Tg(fli1a:EGFP)*^*y1*^ background as described previously (*1*). Embryos imaged at developmental stages later than 36 hpf were treated with 1-phenyl-2-thiourea to inhibit pigment formation (*3*). Zebrafish husbandry and research protocols were reviewed and approved by the NICHD Animal Care and Use Committee.

### Genotyping assays for *cds2*^*y25*^ and *cds2*^*y54*^ mutants

***cds2*^*y25*^**: Zebrafish carrying the *cds2*^*y25*^ mutation were genotyped using KBiosciences Competitive Allele-Specific PCR genotyping system (KASP) assays. Assays were performed on zebrafish adult or embryonic genomic DNA extracts. Primer mixes with two 5’ fluor-labeled oligos that bind to allele-specific primers are provided with a common 3’ oligo, as per the manufactures design. y25FAM, 5’-CAGCGGGAGGAGCC TCTTC-3’ and y25HEX-5’-GCAGCGGGAGGAGCCTCTTT-3’; y25common, 5’-AAATGA AGCGATGGTATTTGCTGAGGATT-3’. Details on assay and primer design can be obtained from KBioscience. The following PCR program was designed to determine wild-type versus mutant PCR fragments: 1) 94^°^C, 15min; 2) 94^°^C, 20s; 3) 61^°^C 1min; 4) GOTO step 2 9x and decrease temperature in step 3 by 0.6^°^C each cycle; 5) 94^°^C, 10s; 6) 55^°^C, 1min; 7) GOTO step 5 35x; 8) 25^°^C, 5min; Plate Read and End. As primers are labeled with fluorophores, homozygous products express only one fluorescent signal while heterozygous products will result in a mixed fluorescent signal.

***cds2*^*y54*^**: PCR was performed on zebrafish adult or embryonic genomic DNA extracts using REDTaq ReadyMix PCR Reaction Mix and the following primers: y54genotypeF-5’-AACAGCTTGATGTAGCACAGCAGAGTA-3’; y54genotypeR-5’-ATGAGGGTGTGGATGATGATGATA-3’. PCR products were then digested with MseI cutting the PCR products carrying mutant *cds2* into 193bp and 124bp fragments versus wild-type *cds2* products that do not cut generating a 317bp PCR product. All fish were housed as pairs in genotyping tanks (R&D Aquatics).

### Zebrafish Expression Constructs and Morpholinos

pCS2(+)-*zvegf*_*165*_ was a gift from Dr. Ruowen Ge (National University of Singapore). CMV promoter-driven zebrafish *vegf*_*165*_ DNA for injection was generated as previously described. DNA was injected at a final concentration of 75 ng/uL.

Morpholino antisense oligonucleotides (Genetools) used in this study include:

*cds2* ATG MO: 5’-TCGCTGTCGTAATTCTGTCATGGTG-3’ that targets −4 to 21 of the 5’ untranslated region and coding region of *cds2* (1.8ng was used as a maximal dose);

*vegfaa*: 5’-CTCGTCTTATTTCCGTGACTGTTTT-3’ (6ng was used as a maximal dose). The *cds2* and *vegfaa* morpholinos used have both been previously validated (*1, 5*). Morpholino injections were performed on zebrafish embryos in the 1-4 cell stage as previously described (*4*). Maximal doses were determined by titration of morpholinos to levels that yielded maximal vascular-specific phenotypes with little to no nonspecific toxic effects.

### siRNA Transfection and Validation

Invitrogen single SilencerSelect or Stealth siRNAs for CDS2 versus a siRNA negative control (control) were purchased and resuspended in H_2_O at a concentration of 10 uM. siPORT Amine (Ambion) was used as a transfection agent (siRNA target sequences in Supplemental Table 1). A double transfection protocol was used as previously described (*6, 7*) with siPORT Amine and a final concentration of 50 nM siRNA per transfection condition. Validation of CDS2 siRNA suppression was previously reported (*1*) but reconfirmed here by western blot. The siRNA sequences utilized are as follows:

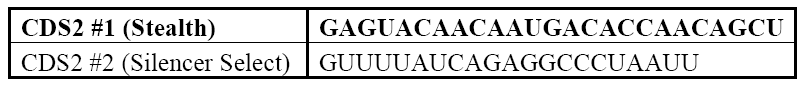

### Endothelial Cell Culture and Assays

HUVECs (Lonza) were cultured in bovine hypothalamus extract, 0.01% Heparin and 20% FBS in M199 base media (Gibco) on 1mg/mL gelatin-coated tissue culture flasks. HUVECs were used from passages 3-6.

3-dimensional (3D) *in vitro* angiogenesis assays were done essentially as described (*6, 7*), 2.5mg/mL collagen type I (BD Bioscience, Acid Extracted) gels were prepared including SCF, SDF1α and IL-3 in the gel at 200 ng/mL and VEGF-A_165_ at the indicated doses described in the manuscript. HUVECs were seeded on the gel surface at a density of 40,000 cells/well. Culture media included FGF and IGF-II. Assays were fixed in 3% glutaraldehyde at the indicated time points and processed for future analysis. Cells were stained with 1% toluidine blue in 30% methanol to increase visualization before imaging. Data analysis was done by imaging the endothelial cell invasive front at 3 depths below the monolayer of cells (approximately 50uM apart in depth). Images were obtained and the number of endothelial cells present at each depth counted; the counts at the 3 depths were added together to give a total number of invading cells per well. A minimum of 5 replicate wells was averaged to give a mean ± SD within a single experiment. At least 2 experimental replicates were performed. Data is represented as a percentage of the 10ng/ml VEGF Control siRNA condition (Figure 1J-% Average number of invading cells) or as a percentage of invasion inhibition at each indicated VEGF dose (Figure 1H-calculated by taking *% Invading number of CDS2 KD cells/% Invading number of Paired Control cells = % Invasion Inhibition).*

### PI(4,5)P2 Quantification

siRNA treated HUVECs were plated on collagen type I-coated plastic wells, allowed to attach overnight and then treated with the indicated doses of rhVEGF-A for predetermined times prior to PIP2 quantification.

PIP2 quantification was done using a competitive ELISA Kit purchased from Echelon Biosciences (Echelon Biosciences, #K-4500) or utilizing a direct ELISA protocol for measuring PIP2 levels as previously modified and described (*8*). PIP2 lipid was extracted following Echelon Biosciences manufacturer recommendations. Absorbance was read using a plate reader at 450 nm. No detergent was used for washes to maintain the PIP2 lipid integrity. Samples were assessed in triplicate, normalized to cell number as assessed by counting 4 individual high powered fields (trypsin treatment of the cells can alter lipid levels and is not advised if it can be avoided), and results given as mean relative PIP2 levels as compared to control untreated cells ± SD for a single experiment or ± SEM when averaging multiple experiments. At least 2-3 experimental replicates were performed.

For measurements of PIP2 levels in lung ECs and tumor associated ECs, competitive ELISAs protocols were done as described above. Results were normalized to input protein levels based on Western blot tubulin expression.

### Injection of PI(4,5)P2 Liposomes

Purified PI(4,5)P2 liposomes (Echelon Biosciences) were incubated in equal molar concentration (40uM) with lipid carrier constructs (Histone H1 Carrier #2, Echelon Biosciences) for 10 minutes at room temperature prior to intra-vascular injection into either the zebrafish or mice. For zebrafish studies, a single 15nl ‘bolus’ injection of BODIPY labeled PIP2 was delivered over the course of 2-3 minutes into the sinus venosus at 48 hpf. Only fish showing substantial and predominantly vascular dispersion of the BODIPY labeled PIP2 were utilized for ISV quantification analysis. Data analysis was done at 72 hpf. For mouse studies, daily retro-orbital injections of PIP liposomes were done versus lipid carrier alone conditions.

### Immunoanalysis

Zebrafish trunk tissue was collected using no. 55 superfine forceps to remove the head and dissect off the yolk ball. Tissue was directly lysed in 2x Laemmli Sample Buffer containing 5% β-ME and a PhosSTOP tablet (Roche), 10 ul per embryo. HUVEC culture lysates were harvested directly into the same lysis buffer described above. Antibodies: p-Erk1/2, T. Erk, p-Akt, T. Akt (Cell Signaling Technologies), Tubulin (Sigma), CDS2 (ProteinTech, 13175-1-AP) and used at 1:1000 in 5% BSA. Secondary HRP-conjugated antibodies were purchased from Santa Cruz or Invitrogen and used at 1:2000 in 5% milk. Quantification of relative band intensity was performed by ImageJ image analysis software, with results shown from at least two independent HUVEC assays, two independent zebrafish clutches, or two-three independent western blots from pooled EC samples collected from a minimum of 2-4 tumors/lungs.

### Tumor Allografts

All animal studies were carried out according to NIH-approved protocols, in compliance with the *Guide for the Care and use of Laboratory Animals*. B16-F10 or LLC tumor cells were maintained in 10% FBS/DMEM growth media. Prior to tumor injection into B6/C57 mice, the mice were shaved to remove hair at the site of tumor growth and facilitate tumor identification and volume measurements. On the day of injection, the tumor cells were trypsinized and resuspended in PBS. 200 ul of PBS/cell suspension containing 10^6^ cells was injected into each flank. For LiCl studies, normal saline (control), LiCl (400 mg/kg), myoinositol (400 mg/kg), or LiCl+myoinositol in 200 uL normal saline were injected daily IP. For vMO (Gene Tools) studies, control vMO (sequence: CCCTGTCCCCTACTTCGCTCATGCT), CDS2 vMO #1 (CCTCTGCCG TAGTTCGGTCATGCT), CDS2 vMO #2 (GGCTTGTCAACGATACTTACAATCA) were injected through the tail vein or retro-orbitally at a dose of 40 ul of 2000 nmol stock per mouse per day. vMO were stored at room temperature and heated at 65^°^C for 15 min prior to use. To verify suppression of the CDS2 gene in vMO treated mice, endothelial cells were isolated from the lungs at the termination of the studies (see below), and lysed for protein analysis to verify protein suppression. L-690,488 chemical inhibitor was purchased from Tocris (#0682), resuspended in DMSO at 100mM and supplied via retro-orbital injection at a daily dose of 80uM per animal daily. For endothelial specific genetic suppression of Cds2, tamoxifen was injected IP daily at a dose of 1mg/mouse in corn oil for 5 days prior to allograft of the tumors. The Cds2 mouse strain used for this research project was created from ES cell clone EPD0033_1_C08 obtained from KOMP Repository (www.komp.org) and generated by the Wellcome Trust Sanger Institute (*9*). Endothelial cells were isolated from both lungs and tumors at the end of the 14 day experiment to confirm Cds2 protein suppression. 8 groups were generated:

Wild type + TMX (no Cre, no floxed Cds2, tamoxifen added CONTROL)

*Cad5*(PAC)-*Cre*ERT2 +TMX (cre driver alone, tamoxifen added CONTROL)

*Cad5*(PAC)-*Cre*ERT2 (cre driver alone, no tamoxifen CONTROL)

*Cds2*^lox/lox^ +TMX (homozygous floxed Cds2, no cre, tamoxifen added CONTROL)

*Cds2*^lox/lox^ (homozygous floxed Cds2, no cre, no tamoxifen CONTROL)

*Cad5*(PAC)-*Cre*ERT2*;Cds2*^lox/+^ +TMX (heterozygous EC-specific knockout)

*Cad5*(PAC)-*Cre*ERT2*;Cds2*^lox/lox^ +TMX (homozygous EC-specific knockout)

*Cad5*(PAC)-*Cre*ERT2*;Cds2*^lox/lox^ (cre driver, homozygous floxed Cds2, no tamoxifen CONTROL)

For all experiments, calipers were used to measure the approximate tumor volume daily, with final weight measurements (mg) made at the termination of the experiment. Tumors were allowed to grow until control tumors reached a volume of approximately 500-1000 mm^3^, day 18 was reached, or significant impairment to the mouse motility from tumor burden was noted.

### Measurement of Tumor Vascularization

B16-F10 or LLC tumors or livers from the treatment mice were collected for analysis and paraffin embedded for H&E and immunofluorescent staining or preserved in OCT medium for cryo-sections. At minimum, 4 livers or 8 tumors of each type were embedded and serially sectioned for vessel density analysis. Immunofluorescent labeling of CD31/PECAM (BD Biosciences #550274, 1:10) and active Caspase3 (Sigma, 1:1000) was done and slides counterstained with HOECHST (Molecular Probes #33342, 1:2000). Images were taken using a Leica Inverted TCS-SP5 II confocal microscope and 20x objective. Three images were taken randomly from within the liver or the tumor body and the average vessel area and average vessel diameter measured per high powered imaging field. Average vessel area was measured using ImageJ, measuring the average CD31/PECAM+ area per high power image field. Average vessel widths were calculated by measuring the distance across the cross-sectioned lumen diameter. 50 measurements were made across vessels per high power image. The data are reported as a mean ± SEM.

### Quantification of Tumor Cell Proliferation

For tumor cell proliferation in the presence of LiCl: B16 or LLC tumor cells were plated in 24 well plates in 500 ul of 10% FBS/DMEM. Cells were allowed to attach and LiCl or vMO added. Estimated concentrations of LiCl were determined proportionally based off of doses provided *in vivo* to the mice (whole mouse dose/approximate tissue volume = x concentration). 10uM (estimated equivalent concentration to mouse dose) and 20uM (estimated double concentration to mouse dose) concentrations of LiCl were added to the growth media daily and proliferation assessed over a multi-day time course. Culture media was changed completely every 3 days for the duration of the time course. For tumor cell proliferation in the presence of vMO: Concentrations for the vMOs were determined based off of the estimated mouse blood volume (approximately 2ml) and proportioned to the volume of tissue culture media present in the assay. A daily dose of 4ul of the 2000nmol stock per well was added to 24 well plates containing 500 ul of 10% FBS/DMEM. Media was changed completely every three days. For both assays, cells were fixed using 2% PFA and stained with HOECHST dye. 5 distinct images per well were assessed and the number of nuclei per well counted. The data are reported as the averages between images ± SD.

### Isolation of mouse lung microvascular endothelial cells or tumor associated endothelial cells

Isolation of mouse lung or tumor microvascular endothelial cells was performed essentially according to a previously described protocol (*10*). Modifications are briefly described, lungs were removed from the mice, washed in 10% FBS/DMEM, minced into 1-2 mm squares and digested with Collagenase Type I (2 mg/ml, Gibco) at 37^°^C for 2 h with occasional agitation. The cellular digest was filtered through a 70 um cell strainer, centrifuged at 1500 rpm and the cells immediately incubated with Dynabead sheep anti-rat IgG (Invitrogen) coated with a rat anti-mouse ICAM-2 mAb (3C4) at room temperature for 10 minutes. The bead-bound cells were recovered with a magnet, washed four times, and the cells immediately collected for protein lysates and PIP2 ELISA protocols. All isolation steps were done in the presence of phosphatase inhibitors.

### Statistics

Statistical analysis of data was done using SPSS or Microsoft Excel. Statistical significance was set at a minimum of P ≤ 0.05, and is indicated in individual figure legends. Student t-tests were used when analyzing 2 groups within a single experiment.

### Study Approval

Zebrafish and mouse husbandry and research protocols were reviewed and approved by the NICHD and NIDCR Animal Care and Use Committee at the National Institutes of Health. All animal studies were carried out according to NIH-approved protocols, in compliance with the *Guide for the Care and use of Laboratory Animals*.

